# A novel virus lineage is abundant in metaviromes from *Dehalococcoides*-containing mixed cultures

**DOI:** 10.64898/2026.09.15.751708

**Authors:** Camilla L. Nesbø, Nadia Morson, Olivia Molenda, Line Lomheim, Julien Lossouarn, Karen L. Maxwell, Elizabeth A. Edwards

## Abstract

*Dehalococcoides mccartyi* are obligately anaerobic organohalide-respiring bacteria that play important roles in the detoxification of chlorinated pollutants in groundwater and sediments. Despite having small genomes, they host a diverse set of mobile elements. Here we characterize a family of mobile elements, termed *integrative and mobilizable element 1* or IME1, comprising 20 from *Dehalococcoides* and one from *Dehalogenimonas alkenigignens*. IME1s are 20,930 – 28,058 bp and are found both integrated in the genomes and as circular episomes. Bioinformatic characterization of IME1 encoded proteins revealed a highly conserved structure with 14 hierarchical orthologous groups (HOGs) found in all 21 IME1s. IME1s lack recognizable hallmark proteins of tailed bacterial viruses (or tailed phages) but encode proteins with similarities to those of filamentous bacterial viruses. In particular, one conserved HOG shows sequence similarity to the pI-like ATPase, the only conserved marker protein identified across filamentous bacterial viruses. Additionally, IME1s encode several small proteins with predicted transmembrane domains and signal peptides, another feature used to identify filamentous bacterial viruses. Both features are also found in budding archaeal viruses of various morphotypes. IME1s dominated metaviromes obtained from the *Dehalococcoides*-containing KB-1 mixed culture and electron micrographs of the corresponding viral fractions revealed abundant filamentous virus-like particles. We therefore propose that the IME1s represent a novel lineage of double stranded budding, likely filamentous, viruses. Database searches suggest IME1s are found in Dehalococcodia and other Chlorofexota but are so far restricted to this phylum.

**Importance:** *Dehalococcoides* are obligate organohalide respiring anaerobic bacteria that use halogenated organic compounds as terminal electron acceptors and are important for bioremediation of polluted sites. Here we describe IME1s, a novel group of viruses associated with Dehalococcoidia that we propose have double stranded DNA (dsDNA) genomes and a non-lytic, likely filamentous lifestyle. IME1s share features with both characterized filamentous bacterial viruses, which package small single stranded DNA (ssDNA) genomes, and archaeal budding viruses, often with larger dsDNA genomes. IME1s expand the known diversity of bacterial viruses and may help reveal how viruses influence the bacteria that detoxify contaminated sites.

## Introduction

Bacteria from the genus *Dehalococcoides* or *Dehalogenimonas,* within the class Dehalococcoidia of the phylum Chloroflexota, are obligate organohalide-respiring bacteria (1) that decontaminate soil and groundwater polluted with chlorinated solvents (2–4). Based on currently available genomes, *Dehalococcoides* fall into three species-level lineages in the genome taxonomy database (GTDB; (5)); *Dehalococcoides mccartyi*, *Dehalococcoides mccartyi*_A and *Dehalococcoides mccartyi*_B, corresponding to previously named Cornell, Victoria and Pinellas groups. These organisms have highly streamlined core genomes of 1.3-1.5 Mb, with multiple key reductive dehalogenase genes concentrated in regions associated with mobile genetic elements (MGEs) (4, 6). This highlights the importance of horizontal gene transfer (HGT) in their adaptation to halogenated pollutants.

We previously investigated *Dehalococcoides* MGEs in an anaerobic mixed culture used for bioremediation, called KB-1 (7). The multiple *Dehalococcoides* strains found in KB-1 subcultures belong to Pinellas and Cornell groups, now classified by GTDB as *Dehalococcoides mccartyi*_B and *Dehalococcoides mccartyi*, respectively. Eight complete fully closed *Dehalococcoides* genomes were assembled from KB-1 subculture metagenomes, revealing a complex interplay of closely related strains participating in reductive dechlorination in this consortium (Supplemental Text S1). Notably, some strains harbour proviruses (prophages) but lack CRISPR-Cas systems, whereas others show the converse pattern. Molenda *et al*., (7) characterized an integrative and mobilizable element (IME) carrying the *vcrA* gene, which encodes the vinyl chloride reductase responsible for the transformation of vinyl chloride to non-toxic ethene. This VcrA-IME was targeted by the strain’s CRISPR-Cas system and was found both integrated into *Dehalococcoides* genomes and as a circular episome. The same study identified a second family of larger mobile elements, termed IME1s, that were similarly targeted by CRISPR-Cas and found both integrated and as circularized episomes. The ∼20kb IME1s were larger than the VcrA-IME (∼12kb), did not carry known *rdhA* genes (7), and their biological function remained unknown.

Roux *et al.*, (8) used machine learning to identify previously unrecognized viruses related to the Inoviridae, which include most characterized filamentous bacterial viruses (filamentous phages). They found that filamentous viral sequences were much more pervasive and diverse than had been previously considered among prokaryote (meta)genomes. Notably, their analysis classified two KB-1 metagenomic contigs corresponding to the IME1 sequences as members of a proposed family of putative filamentous viruses within the Inoviridae, the Densinoviridae (sub family Sf_3). They also identified a full-length IME1 from *Dehalococcoides mccartyi* CG3 and partial IME1 sequences from other *Dehalococcoides* genomes (8). Their pipeline identified potential filamentous viruses based on the presence of the pI morphogenesis protein, the only conserved protein shared across characterized inoviruses, together with genomic features including short genes encoding hydrophobic proteins and small genome size. The pI protein belongs to the FtsK family of P-loop NTPases, and is frequently automatically annotated as Zot (zonula occludens toxin) because it contains a Zot-domain (PF05707;). However, pI functions in virion assembly and extrusion at the bacterial membrane (10, 11). Filamentous bacterial viruses establish non-lytic infections in which virions assemble and bud from the bacterial membrane without killing the host (9). Whether integrated into the chromosome or maintained extra chromosomally, all characterized filamentous bacterial viruses replicate through circular dsDNA intermediates but package ssDNA genomes into the virions.

Here we characterize IME1 sequences and present evidence that they represent a novel virus lineage sharing features with both filamentous bacterial viruses and archaeal budding viruses. We show that IME1 sequences dominate metaviromes prepared from KB-1 cultures collected when the microbial community is actively dechlorinating trichloroethene or vinyl chloride (*i.e.,* optimal growth conditions for *Dehalococcoides*). Electron microscopy of these viral fractions revealed abundant filamentous virus-like particles. Together, these observations support IME1s as a novel lineage of bacterial viruses with a likely non-lytic, filamentous mode of propagation.

## Results and Discussion

### Five new IME1s identified in Dehalococcoides and one in Dehalogenimonas alkenigignens

Based on the genomic organization of the IME1s described in Molenda *et al.*, (7), we identified five new IME1s in *Dehalococcoides* and one new IME1 in *Dehalogenimonas alkenigignens* strain BRE15M (Table 1, Figure 1). The elements range in size from 21 - 28kb (Table 1), *Dehalococcoides* IME1s have a GC content in the range 44 - 45%, compared with 47 - 49% for their host genomes, while the *D. alkenigignens* IME1 has a GC content of 52%, compared with 56% for its host genome. The consistently lower GC content of IME1s relative to their host genomes is consistent with a history of horizontal transfer of these elements.

**Figure 1.**
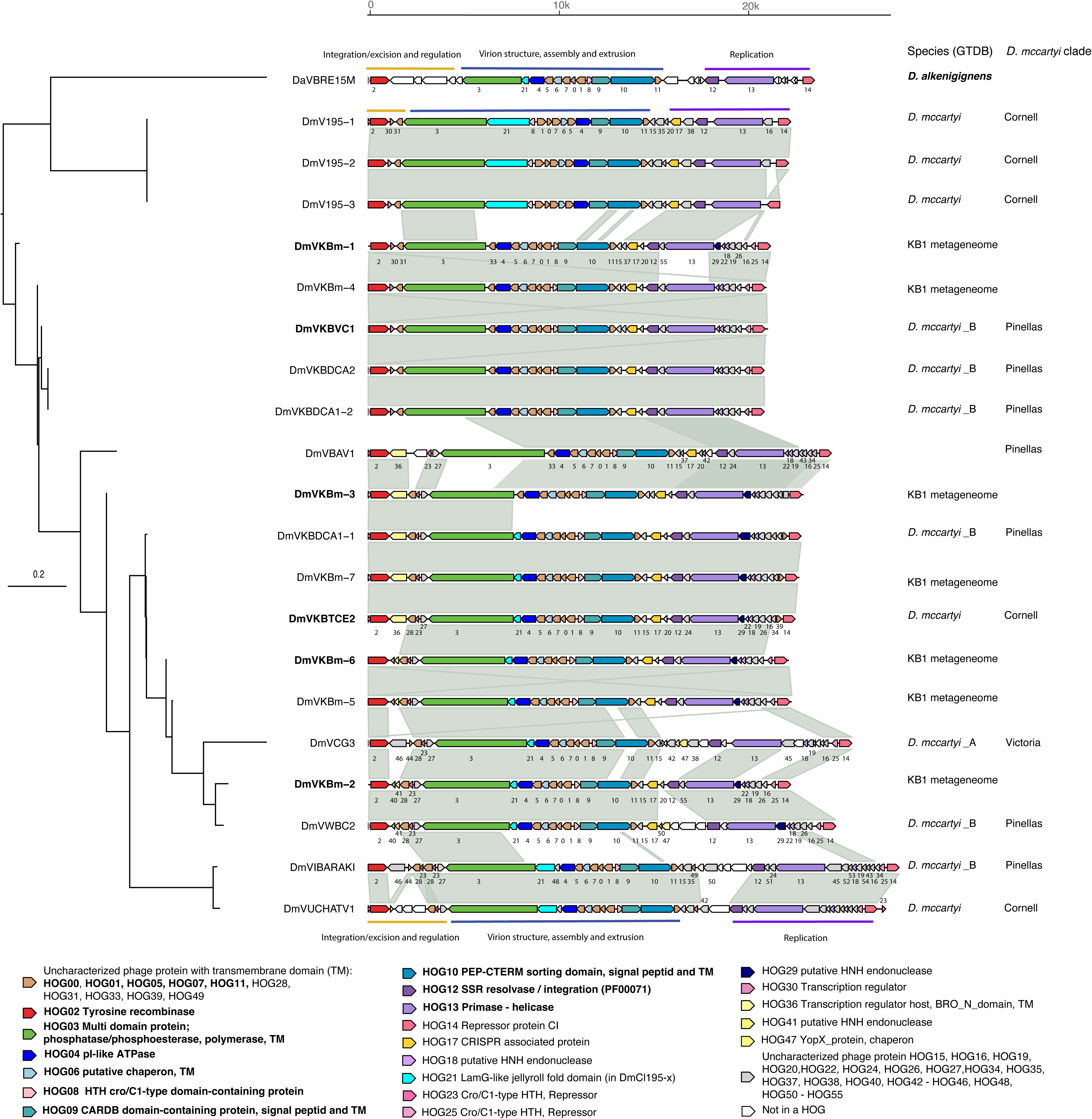
Comparison of IME1 elements identified in Dehalococcoidia genomes and KB-1 metagenomes drawn using gggenomes v1.0.1 in R. The tree on the left is the ‘species’-tree obtained from Orthofinder v3.0.1b1. The sequences were aligned by minimap2 v.2.28. Each CDS is colored according to the HOG it belongs to and numbers under CDSs correspond to HOG number. CDS with no HOG are shown in white. The putative module the CDSs belong to are indicated as horizontal lines. Vertical blocks between sequences indicate sequence similarity. Annotation of the HOGs is shown on the right, and the ones in bold are found in all IME1s. The *Dehalococcoides* clade and/or GTDB species assignment of host genomes are indicated for elements obtained from genome sequences; *Dehalococcoides mccartyi* (*D. mccartyi* / Cornell), *Dehalococcoides mccartyi*_A (*D. mccartyi* A / Victoria), *Dehalococcoides mccartyi*_B (*D. mccartyi* B / Pinellas) and *Dehalogenimonas alkenigignens* (*D. alkenigignens*). Note that because the CP000688 (Strain BAV1) was suppressed in GenBank the IME1 DmVBAV1 does not have a GTDB taxonomy assignment.

**Table 1.** IME1s of *Dehalococcoides mccartyi* (*Dhc*), *Dehalococcoides mccartyi*_A (*DhcA*) *Dehalococcoides mccartyi*_B (*DhcB*) and *Dehalogenimonas alkenigignens* (*Dhg*) as Dehalococcoidia Chromosomal Islands (DeCIs). IME1-1 – IME1-14 were previously identified by Molenda *et al*. (7).

| Name | Molenda et al. 2019 | Attachment site (att<sub>c</sub>) | Length (Kb) | Host genome | Accession number | Dhc Clade | GTDB taxonomy | %GC IME1 (genome) |
| --- | --- | --- | --- | --- | --- | --- | --- | --- |
| DmVKBm-1 | IME1-1 |  | 21.3 | *Dhc* KB-1 metagenome | PRJNA376155<sup>a</sup> |  |  | 44.4 |
| DmVKBm-2 | IME1-2 |  | 22.3 | *Dhc* KB-1 metagenome | PRJNA376155<sup>a</sup> |  |  | 44.6 |
| DmVKBm-3 | IME1-3 |  | 23.0 | *Dhc* KB-1 metagenome | PRJNA376155<sup>a</sup> |  |  | 44.6 |
| DmVKBm-4 | IME1-4 |  | 21.0 | *Dhc* KB-1 metagenome | PRJNA376155<sup>a</sup> |  |  | 44.7 |
| DmVKBm-5 | IME1-5 |  | 22.4 | *Dhc* KB-1 metagenome | PRJNA376155<sup>a</sup> |  |  | 44.5 |
| DmVKBm-6 | IME1-6 |  | 22.2 | *Dhc* KB-1 metagenome | PRJNA376155<sup>a</sup> |  |  | 44.4 |
| DmVKBm-7 | IME1-7 |  | 22.8 | *Dhc* KB-1 metagenome | PRJNA376155<sup>a</sup> |  |  | 44.5 |
| DmVKBTCE2 | IME1-8 | tRNA-Ala | 22.5 | *Dhc* strain KBTCE2 | CP019865 | Cornell | *Dhc* | 44.4 (49.1) |
| DmCKBVC1 | IME1-9 | tRNA-Ile | 21.9 | *Dhc* strain KBVC1 | CP019968 | Pinellas | *DhcB* | 44.7 (47.3) |
| DmCKBDCA1-1 | IME1-10 | tRNA-Ala | 22.9 | *Dhc* strain KBDCA1 | CP019867 | Pinellas | *DhcB* | 44.6 (47.4) |
| DmVKBDCA1-2 | IME1-11 | tRNA-Ile | 20.9 | *Dhc* strain KBDCA1 | CP019867 | Pinellas | *DhcB* | 44.8 (47.4) |
| DmVKBDCA2 | IME1-12 | tRNA-Ile | 20.9 | *Dhc* strain KBDCA2 | CP019868 | Pinellas | *DhcB* | 44.8 (47.3) |
| DmVWBC2 | IME1-13 | tRNA-Ile | 24.7 | *Dhc* strain WBC-2 | CP017572 | Pinellas | *DhcB* | 45.1 (47.4) |
| DmVBAV1 | IME1-14 | tRNA-Ala | 24.4 | *Dhc* strain BAV1 | CP000688 | Pinellas | NA<sup>b</sup> | 43.4 (47.2) |
| DmV195-1 |  | tRNA-Lys | 22.2 | *Dhc* strain 195 | CP000027 | Cornell | *Dhc* | 45.8 (48.9) |
| DmV195-2 |  |  | 22.2 | *Dhc* strain 195 | CP000027 | Cornell | *Dhc* | 45.7 (48.9) |
| DmV195-3 |  |  | 22.1 | *Dhc* strain 195 | CP000027 | Cornell | *Dhc* | 45.8 (48.9) |
| DmVCG3 |  | tRNA-Ile | 25.5 | *Dhc* strain CG3 | CP013074 | Victoria | *DhcA* | 44.3 (46.9) |
| DmVIBARAKI |  |  | 28.0 | *Dhc* strain IBARAKI | AP014563 | Pinellas | *DhcB* | 43.6 (47.0) |
| DmVUCHATV1 |  | tRNA-Ala | 27.3 | *Dhc* strain UCH-ATV1 | AP017649 | Cornell | *Dhc* | 43.8 (48.8) |
| DaVBRE15M |  | tRNA-Ile | 22.9 | *Dhg* strain BRE15M | QEFQ01 | ---- | *Dhg* | 52.1 (56.3) |

a) Metagenomic reads and MAG assemblies are available under BioProject PRJNA376155.
b) CP000688 is suppressed in GenBank and therefore has no associated GTDB taxonomy.

IME1 were found in members of all three *Dehalococcoides* GTDB species (Figure 1), although they were not present in all genomes examined. Of the 24 fully closed genomes examined, 10 contained at least one IME1. The 21 IME1s were renamed using the first letters of the host genus and species, followed by V for virus and the strain designation; “KBm” denotes elements identified in KB-1 metagenomic contigs that could not be linked to a strain. Some *Dehalococcoides* genomes contained multiple IME1s. *D. mccartyi*_B strain KBDCA1 contains two divergent IME1s, DmVKBDCA1-1 and DmVKBDCA1-2, whereas the three IME1s in *D. mccartyi* 195 are almost identical (Figure 1).

### IME1s encode features of filamentous and budding viruses

Predicted IME1 coding sequences (CDSs) were clustered into 56 hierarchical orthologous groups (HOGs) (Table S1), with most CDSs assigned to a HOG (Figure 1). Fourteen HOGs were conserved across all 21 IME1s analysed, with one additional HOG conserved across all *Dehalococcoides* IME1s. Note that the HOG designations used here are specific to this dataset. A phylogenetic tree constructed from the concatenated HOG alignments revealed a high degree of conservation among *Dehalococcoides* IME1s (Figure 1). Notably, this phylogeny does not match the GTDB-genome taxonomy or their *Dehalococcoides-*clade designation (Figure 1). This, together with the variable distribution of the IME1s, suggest extensive transfer and loss of these elements.

IME1s lacked recognizable homologues of previously described *Dehalococcoides* tailed virus proteins and hallmark proteins of tailed viruses (Caudoviricetes). However, many proteins had best structural matches to other proteins in viral databases (BFVD 2023_2 and PHROG; Table S2.1–Table S2.21) that are not part of the hallmark set for tailed viruses. Comparative analysis revealed a conserved architecture comprising three major regions associated with putative (i) integration/excision and regulation, (ii) virion biogenesis (i.e. structure, assembly and extrusion), and (iii) replication (Figure 1).

The conserved central region of IME1s contains two features characteristic of filamentous bacterial viruses: a pI-like protein and multiple short hydrophobic proteins with predicted transmembrane helices. Related ATPases and hydrophobic structural proteins are also found in several budding viruses (32–34). Both features are encoded within the putative virion biogenesis module of IME1s (Figure 1). Together, these observations support the proposal by Roux et al. (8), that IME1s represent a novel lineage of filamentous and/or budding viruses.

Specifically, HOG04 proteins show both sequence and structural similarity to pI-like proteins from filamentous viruses and to archaeal DNA-packaging ATPases (Table S2), the same protein family used by Roux *et al*. (8) to identify cryptic inoviruses. A phylogenetic tree constructed from a structural alignment of HOG04 proteins and their best structural matches placed them as a distinct lineage in a clade that also includes pI-proteins from characterized inoviruses, as well as metagenomic sequences, (Figure S1). The pI-like proteins are ATPases involved in DNA packaging, virion assembly, and filamentous virion extrusion (8, 9, 15), and we hypothesize that HOG04 performs a similar function in IME1s.

Immediately downstream of HOG04, all IME1s encode five conserved HOGs (HOG05–HOG07, HOG00, HOG01) containing predicted transmembrane domains, with some also containing predicted signal peptides (Table S2). This region is inverted in the three IME1s from strain 195 (DmV195-1–3). The best structural matches of HOG00 proteins were to the VP1_SSV1 structural protein from *Sulfolobus* spindle-shaped virus 1 (SSV1) and phrog_11172 (containing the VP1_SSV1 gene) in the Prokaryotic Virus Remote Homologous Groups database (16) (Table S2). Several HOG01 proteins also showed structural similarity to these proteins. SSV1 VP1 is a hydrophobic structural protein that localizes to the membrane interface, participates in budding, and is the major capsid protein of this virus. We hypothesise that HOG00 and HOG01 may perform related functions during IME1 particle assembly and release. Some HOG00 and HOG01 proteins also showed structural similarity to uncharacterized proteins from tailed bacterial viruses, although these matches were weaker than those to phrog_11172-proteins.

A lambda repressor-like protein (HOG08), encoded on the opposite strand, may regulate expression of the putative virion genes. Three additional conserved genes (HOG09–HOG11) encode proteins with predicted transmembrane and signal peptides, several of which also show structural similarity to viral proteins (Table S2). HOG09 and HOG10 contain CARDB and PEP-CTERM domains, respectively, which are associated with cell adhesion and membrane localisation. Although gene content and synteny are highly conserved across this region, several HOGs show relatively low sequence conservation (Figure 1).

The two modules flanking the conserved central module are more variable in gene content. In the ‘integration / excision and regulation module’, the conserved tyrosine recombinase HOG02, located at the 5′ end of all IME1s, likely mediates integration into and excision from the host chromosome (9). Phylogenetic analysis revealed that HOG02 homologues are widely distributed among Chloroflexota and are not unique to IME1, suggesting extensive horizontal gene transfer of these genes (Figure S2). HOG02 is followed by two to five variable CDSs representing nine HOGs, including transcriptional regulators (HOG23 or HOG30) and proteins associated with DNA binding and DNA metabolism (e.g. Bro-N domain proteins and HNH endonucleases). Several IME1s also carry putative accessory genes in this region; for example, DaCIBRE15M encodes two proteins with similarity to restriction endonucleases.

The 3’ replication module is also variable in gene content but contains three highly conserved HOGs present in all IME1s. HOG12 encodes an SSR resolvase, which may be involved in replication and excision (9) and/or maintenance of the episomal state (17). HOG13 encodes an archaeal primase–helicase associated with bacterial and archaeal viruses and at the 3′ end, all IME1s encode a repressor (HOG14). HOG17, present in 17 IME1s, shows similarity to the CRISPR-associated protein Cmr6/4, although its function remains unclear. Other proteins with predicted functions in this module are also associated with DNA metabolism and regulation, including HNH endonucleases (HOG18 and HOG19), a Cro/C1-type repressor (HOG25), and a YopX-like chaperone (HOG47).

### IME1s are highly abundant in KB-1 metaviromes

To investigate if IME1s and other MGEs are present in extracellular viral fractions, we collected viral fractions from two KB-1 mixed microbial cultures: the first grown with trichloroethene as electron acceptor (referred to as the TCE-virome) and the second a subculture grown on vinyl chloride (VC-virome). These cultures are referred to as KB-1/TCE-MeOH and KB-1/VC-MeOH, as methanol (MeOH) was supplied as electron donor to each mixed culture. Previous metagenomic analyses detected circular IME1s in these cultures, although their abundance varied among samples (7).

DNA extracted from each viral fraction was used to construct two metaviromes (one from each culture) which were analysed to identify MGEs, including bacterial viruses and IMEs, and to quantify their relative abundances. Of the ∼70% of Illumina reads that could be assigned a taxonomy, >80% were classified as *Dehalococcoides*, despite *Dehalococcoides* typically comprising ∼40% of the KB-1 community based on 16S rRNA abundance.

When metavirome reads were mapped to the five previously closed KB-1 *Dehalococcoides* genomes from the TCE- and VC-grown cultures, IME1s were by far the most highly enriched regions in both datasets (Figures 2 and S3; Table 2). In the TCE-metavirome, 79% of Illumina reads mapped to IME1 sequences. For the VC-metavirome 56% of the reads mapped to IME1s (Table S3). IME1 regions reached read depths of ∼39,000x – 56,000x when reads were mapped to *Dehalococcoides* genomes (Table 2, Table S3), whereas known tailed proviruses showed coverage similar to chromosomal background (Table S4). The same enrichment was observed with PacBio CCS reads to *Dehalococcoides* genomes (Table 2) and in an independently prepared TCE virome collected in 2026 (Figure S4).

**Figure 2.**
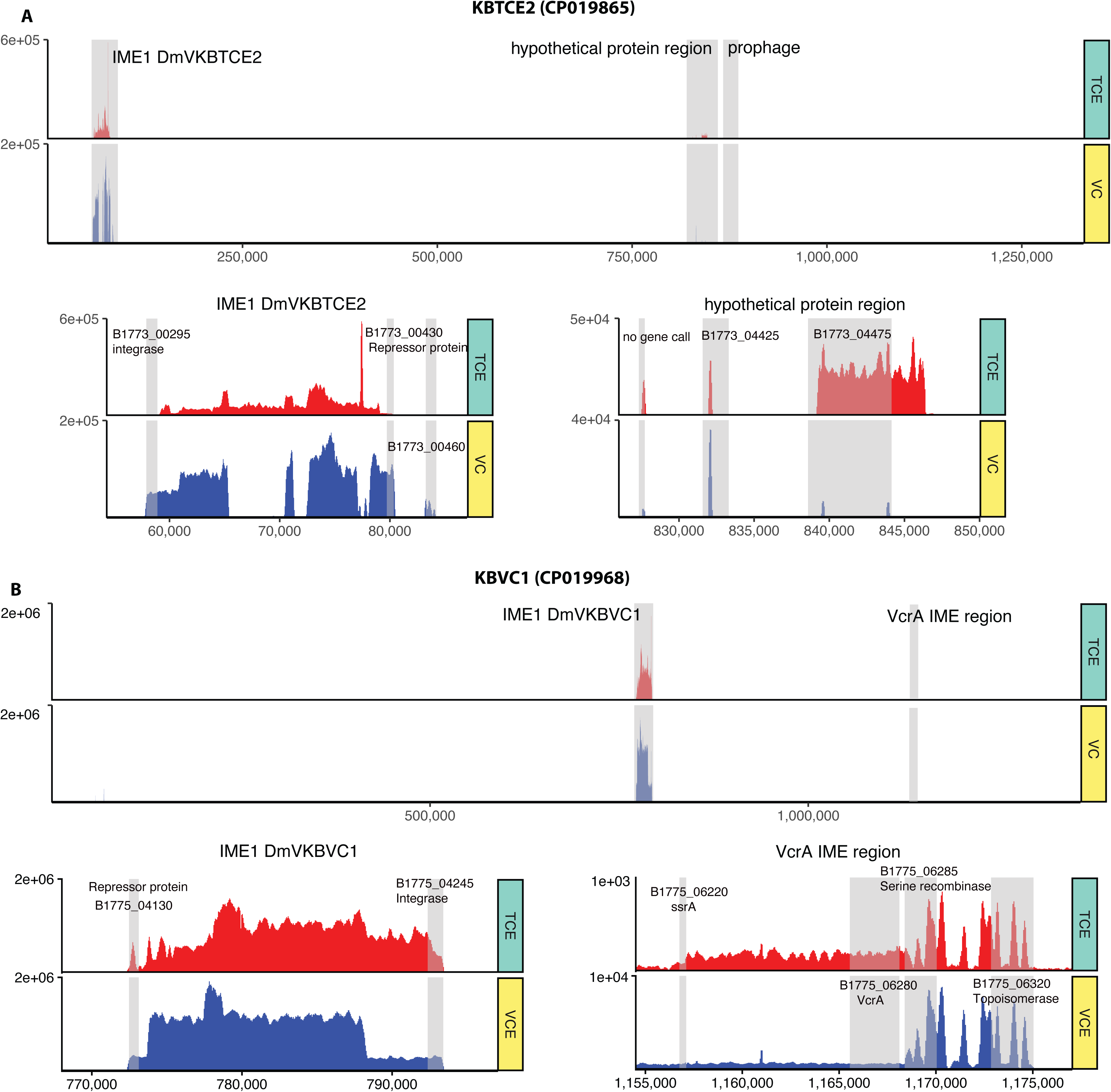
Mapping deduplicated metavirome reads to the *D. mccartyi* KBTCE2 (A) and *D. mccartyi* KBVC1 (B) genomes. The upper panel for each genome shows mapping along the full genome sequence, with read depth shown on the Y-axis for the TCE metavirome (TCE) and VC metavirome (VC). Regions showing high read recruitment are zoomed in and highlighted below for each genome. Note that both viromes contain genomic *D. mccartyi* sequences. The average read-depth to the KBTCE2 (CP019865) genome, excluding the IME1 region, was 130x for the TCE-virome and 170x for the VC-virome (Table 2). The average read-depth to the KBVC1 (CP019968) genome, excluding the IME1 region, was 60x for the TCE-virome and 400x for the VC-virome (Table 2). The figure was drawn using ggcoverage v. 1.3.0 in R.

**Table 2.** Average read depth for genomes in Figure 2 and Figure S3. Read depth was calculated using BBmap and samtools’ bedcov for Illumina data and minmap2 in Geneious Prime 2026 for Pacbio CCS reads.

| Genome / Region | Illumina 2019 virome |  | PacBio CCS virome |  | Illumina 2026 virome |
| --- | --- | --- | --- | --- | --- |
|  | TCE virome read depth | VC virome read depth | TCE virome read depth | VC virome read depth | TCE virome read depth |
| **CP019865_KBTCE2** |  |  |  |  |  |
| 1 - 57897 | 61 | 315 | 0 | 0.1 | 91 |
| **57898 – 80347; IME1** | **43282** | **39839** | **929** | **1170** | **16459** |
| 80348 - 1329198 | 131 | 214 | 1 | 0.1 | 82 |
| chromosome without IME1 | 128 | 219 | 1 | 0.1 | 82 |
| **CP019968_KBVC1** |  |  |  |  |  |
| 1- 772474 | 74 | 516 | 0.1 | 0.5 | 74 |
| **772475 – 793406; IME1** | **56944** | **51867** | **1101** | **1233** | **11392** |
| 1156755 – 1168114; VcrA-IME | 139 | 482 | 0 | 1.3 | 66 |
| 793406 - 1359904 | 33 | 482 | 0.1 | 0.3 | 63 |
| chromosome without IME1 | 57 | 409 | 0.1 | 0.3 | 70 |
| **CP019999_KBTCE1** |  |  |  |  |  |
| 1 - 1388914 | 30 | 324 | 0.1 | 0.8 | 67 |
| 1235442 -1250194; VcrA IME | 271 | 1468 | 3.5 | 6.1 | 91 |
| **CP019969_KBVC2** |  |  |  |  |  |
| 1 - 1337731 | 30 | 321 | 0.1 | 0.8 | 70 |
| 1185817 – 1210000; VcrA IME | 175 | 1042 | 2.2 | 4.4 | 71 |
| **CP019866_KBTCE3** |  |  |  |  |  |
| 1 - 1271604 | 103 | 188 | 1 | 0.6 | 83 |

While metavirome reads also mapped to genomic regions of *Dehalococcoides*, particularly for the VC-metavirome (Figure 2, Supplemental Figure 1, Table 2) the background read depth was very low compared to that of IME1s. Specifically, background read depth was 30x - 130x for the TCE-virome reads mapped to *Dehalococcoides* genomes and in the range 170x - 400x for VC-virome reads mapped to *Dehalococcoides* genomes (Table 2). This background likely represents cellular contamination of the viral fraction, as the small size of *Dehalococcoides* may allow some cells to pass through the 0.45 µm filter (Figure 3). Because the viral fractions were treated with DNase I before particle disruption and DNA extraction, the strong enrichment of IME1 sequences is consistent with protection of IME1 DNA within extracellular particles.

**Figure 3.**
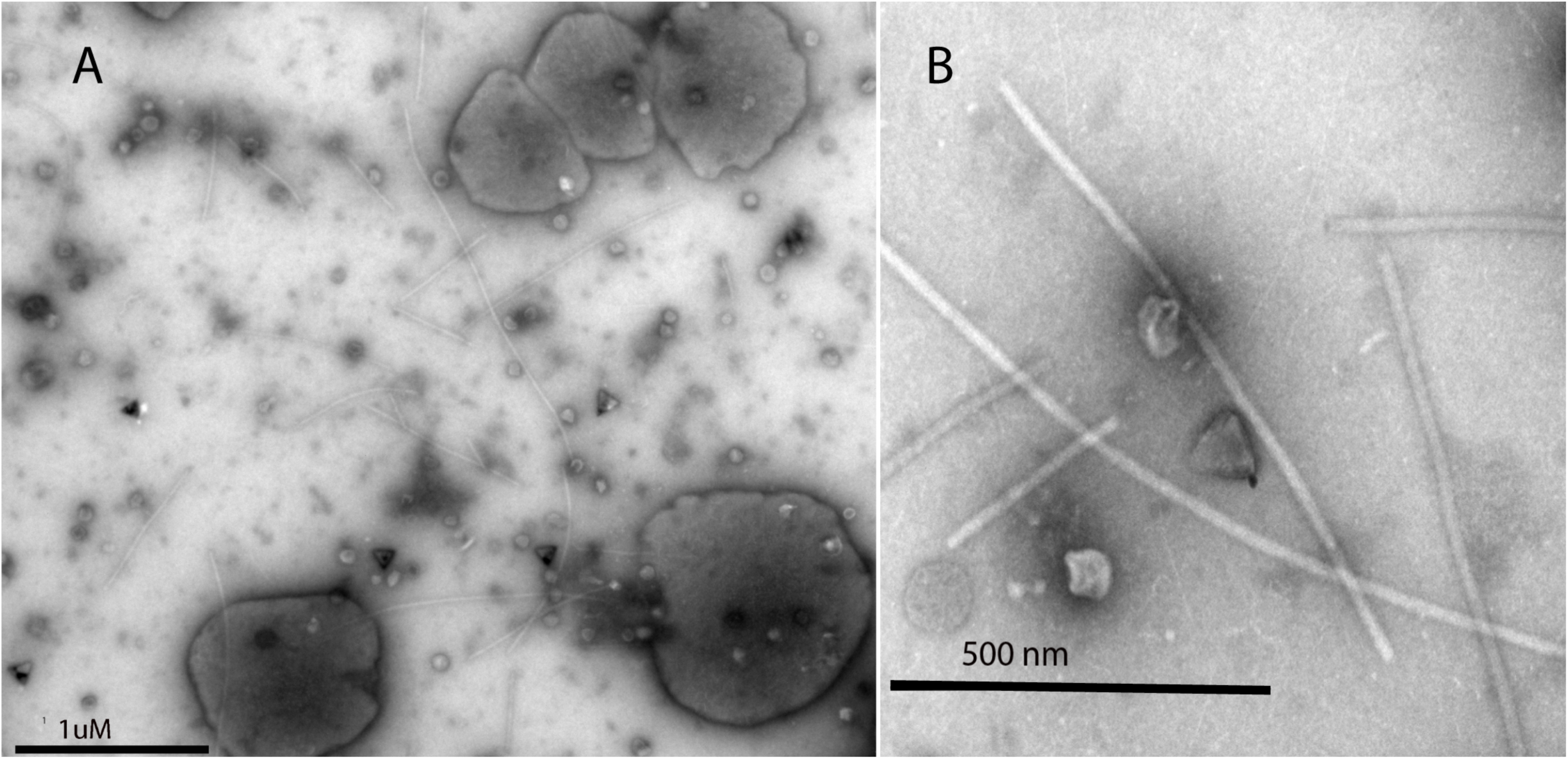
Negatively stained transmission electron microscopy of the KB-1/TCE-MeOH and KB-1/VC-MeOH viral fractions. A: Overview image of the crude virus fraction preparations. The large structures match in size to *Dehalococcoides* cells. Filaments and vesicles are abundant, as well as possible tail-less bacterial-virus-capsids. B: Image focusing on filaments of varying length, round vesicles and possible bacterial virus capsids.

The IME1 populations differed between the two cultures (Table S3). In the TCE-metavirome, DmVKBm-1 was the most prominent element, reaching a mean depth of 11,507×, while DmVKBm-3 dominated the VC-metavirome at 12,026×. DmVKBVC1 was abundant in both viromes (∼10,000×). However, the high sequence similarity among many of the KB-1 IME1s complicates accurate abundance estimates for individual IME1s (Figure 1).

To better resolve the IME1 composition in each metavirome, we applied strict read mapping of Illumina reads to representative KB-1 IME1 sequences, allowing only 1% mismatch and requiring paired reads to map in proximity (Table S3). This analysis identified DmVKBVC1 and DmVKBm-3 as the only previously-assembled-IME1s present at high abundance in the VC-metavirome. Notably, the DmVKBm-3 variant in the VC-metavirome appeared to lack three CDSs in the 3’ region (CDSs 25–27). The TCE-metavirome contained greater IME1 diversity, with DmVKBVC1 and DmVKBm-1 dominant, and DmVKBm-3 also detected. Mapping of the PacBio CCS metavirome-reads supported these results (Table S3). In the independently collected 2026 TCE virome, DmVKBm-3 recruited the greatest number of reads, indicating a shift in the dominant IME1 population over time (Table S3).

The high abundance of IME1s was also evident in in hybrid assemblies combining Illumina and PacBio reads (Table S5). Of 984 TCE-metavirome contigs with >100× sequencing depth, 921 (93%) showed BLASTn matches to IME1 sequences (Table S6). Similarly, 214 of 266 (80%) VC-metavirome contigs with >100× coverage matched IME1s (Table S7). No full-length IME1 sequences were recovered from the TCE hybrid assembly, potentially reflecting the greater IME1 diversity in this culture. No previously described *Dehalococcoides* viruses were detected among the high coverage TCE contigs, although a contig corresponding to the VcrA-IME was identified (Table S6). The VC-metavirome assembly contained longer IME1-matching contigs, as well as contigs assigned to tailed viruses (Table S7). VirSorter2 and VIBRANT identified additional viral contigs in both metaviromes, but these occurred at substantially lower abundance than the IME1 contigs (Tables S8 and S9; Supplemental Text S2). The only viral contigs with read depths > 1000x in either dataset corresponded to IME1 sequences (Table S9).

To improve IME1 assembly, additional assemblies were performed using PacBio reads alone or in combination with 10% of deduplicated Illumina reads. These approaches recovered full-length circular IME1 sequences of 21,059bp (from the VC-metavirome) and 20,995bp (from the TCE-metavirome) with > 99% nucleotide identity to IME1 DmVKBVC1 (Figure S5) from both metaviromes. No full-length tailed bacterial virus genomes were recovered from either metavirome.

### KB-1 viral fractions contain abundant filaments and vesicles

Viral fractions from KB-1 cultures that were prepared for sequencing were also visualized using negatively stained transmission electron microscopy (TEM) (Figure 3). TEM revealed numerous long filamentous structures as well as round vesicles, alongside small cells consistent with the size of *Dehalococcoides* (Figure 3). The abundance of filamentous structures is consistent with the hypothesis that IME1s produce filamentous virions, although the IME1 genome could not be directly linked to a specific particle morphology. Because sonication could potentially damage viral particles, we performed a second viral-fraction preparation without sonication (Supplemental Text S3). This preparation yielded more virus-like particles (VLPs) with recognizable morphologies (Figure S6), while filamentous structures and vesicles remained abundant (Figure S7). Filamentous structures and vesicles have also previously been observed in association with *Dehalococcoides* cells (18).

### IME1 share features with bacterial and archaeal viruses

IME1s share features with both filamentous bacterial viruses, including members of the Inoviridae (9), and with archaeal viruses that egress via budding such as Fuselloviridae (19, 20). The conserved HOG04 proteins, that shows significant similarity to the Inoviridae-hallmark pI-protein also exhibits structural similarity the genome-packaging ATPases from the eukaryotic and archaeal viruses (Figure S1)(20). Another shared feature of filamentous bacterial viruses and archaeal budding viruses is the presence of multiple short, hydrophobic structural proteins, often containing predicted transmembrane helices, that function as major or minor capsid components (8, 9, 12–14). In IME1 we suggest HOG00 (and HOG01) carry out this function, and as mentioned above, HOG00 proteins show structural similarity to the capsid protein from the archaeal virus SSV1.

Both inoviruses and many archaeal viruses (e.g. SSV1) establish non-lytic infections in which virions are released by budding or extrusion without host-cell lysis. The high abundance of IME1 sequences in viral fractions collected during active *Dehalococcoides* growth is consistent with particle production without widespread host lysis and raises the possibility that IME1s similarly establish chronic infections. By contrast, tailed bacterial viruses in the KB-1 system were previously observed to be induced under starvation conditions (21).

All characterized bacterial filamentous viruses possess relatively small genomes (from ∼4 to ∼11 kb) that are packaged as circular ssDNA (9, 11). By contrast, archaeal budding viruses such as Fuselloviridae (19, 20) and archaeal filamentous lytic viruses (recently assigned to the realm Adnaviria (22)) typically possess dsDNA genomes that can be substantially larger ( ∼14 to ∼41 kb for the adnavirians)). IME1 genomes exceed 20 kb and occur as dsDNA when integrated into host chromosomes or maintained as circular episomes. Their high recovery from libraries prepared using dsDNA-adapter ligation is consistent with a dsDNA form in the viral fractions, although the nucleic-acid state packaged within IME1 particles has not been directly determined. Thus, IME1s differ substantially from characterized bacterial filamentous viruses in genome size and may also differ in packaged genome type. Together, these observations support IME1s as a novel lineage of bacterial viruses with features of both filamentous bacterial viruses and archaeal viruses.

Molenda *et al*., (7) proposed that IME1s might be virus satellites that require a helper virus for propagation, while Morson (23) subsequently suggested that they may represent phage-inducible chromosomal islands (PICIs). These interpretations were based largely on the absence of hallmark genes associated with tailed bacterial viruses, and before we recognized the filamentous virus-like features of IME1s. Several observations now argue against a requirement for a helper virus. Most importantly, IME1s encode a conserved candidate virion biogenesis module, including a pI-like assembly/extrusion ATPase and multiple candidate structural proteins, providing a plausible mechanism for particle formation independent of a tailed helper virus. In addition, the two dominant *Dehalococcoides* genomes recovered from the KB-1/VC-MeOH culture showed no consistent association between IME1 and known tailed proviruses: KBVC1 contains an IME1 but no known tailed provirus, while KBVC2 contains a tailed provirus but no IME1 (7). Analysis of *Dehalococcoides* genomes using Satellite finder (24) also failed to identify the IME1s as possible PICIs. While induction of a provirus is not necessarily a prerequisite for piracy, and IME1 could also take advantage of infection by an exogenous tailed virus, the end-result would (in both case) be the lysis of the host caused by the tailed helper virus. In contrast, we did not see evidence of host lysis and massive release of head and tail-like virions. However, we cannot rule out the possibility that IME1s depend on an unidentified lytic helper virus. Some mobile elements can hijack helper-virus particles so efficiently that very few helper virus genomes are detected in the virions produced (e.g (25)). Studying the expression of IME1 in pure *Dehalococcoides* cultures with no possible helper virus would help resolve this, as well as help link the IME1 genomes to their virion particle.

### Other *Dehalococcoides* MGEs and genes in the metaviromes

Mapping metavirome reads to *Dehalococcoides* genomes did not provide strong evidence that other MGEs or *rdh*-containing regions were packaged into virions (Supplemental Text 4; Table S10). The VcrA-IME showed modest enrichment above chromosomal background, but at levels far below those observed for IME1s. We did, however, identify other genes and regions with high read recruitment. For example, B1773_04475 in *D. mccartyi* KBTCE2 (CP019865, Figure 2) showed BLAST similarity to HOG03 in IME1 (65% protein identity to DmVBAV1 CDS 6), suggesting it may represent a remnant of an IME1 or an IME1-like mobile element. Similarly, one of the high-coverage regions in the *D. mccartyi* KBTCE3 (CP019866; B1774_04120-B1774_04140, Figure S1) genome also bears hallmarks of an IME1 remnant, with similarities to HOG03, HOG27, HOG23, and HOG28. The tyrosine-type recombinase/integrase (B1774_00320) in this genome with high read depth (Figure S1), shows low similarity to the integrase in IME1. It is located 4kb upstream of reductive dehalogenase TceA (26). However, only the recombinase shows elevated read depth, suggesting it is part of another mobile element.

### IME1-like elements are found in other Chloroflexota genomes

Examination of sequences classified as *Densinoviridae* Sf_3 by Roux *et al.*, (8) identified an IME1-like element in a Chloroflexi bacterium RBG_13_48_17 MAG (MGMO00000000.1) carrying homologs of eight conserved IME1 HOGs (Table S11). More distantly related IME1-like sequences were identified on short metagenomic contigs for which host assignment was not possible (Table S11).

BLAST searches and phylogenetic analyses based on the pI-like protein HOG4 identified additional IME1-like elements across Dehalococcoidia and Chloroflexota. Six additional complete or partial IME1s were identified in *Dehalococcoides* genomes deposited after the initial set of 21 elements analysed here was defined (Table S12, Figure S8). We also identified candidate IME1-like elements containing homologs of 7–12 IME1 HOGs in five Dehalococcoidia or Chloroflexi MAGs (Table S11, Figures S9). A circular IME1 element was also identified in a Candidatus *Omnitrophota* MAG (JBAZEP010000021; Table S11), however, closer inspection suggested the contig was likely mis-binned and originated from an organism closely related to *Dehalococcoides*. Similarly, short IME1-like contigs assigned to Candidatus Pacearchaeota and Phycisphaerae MAGs may represent mis-binned sequences originating from Chloroflexota (Table S11). Such mis-binning and assembly fragmentation are not unexpected because IME1s can occur in metagenomes both as chromosomal insertions and as independent circular replicons.

### Concluding remarks

Our analyses strongly suggest that IME1s represent a novel virus lineage that share features with both bacterial and archaeal viruses (Table 3). IME1 sequences dominated viral fractions from actively growing *Dehalococcoides*-containing cultures, reaching abundances orders of magnitude above chromosomal background and known tailed bacterial viruses. Together with the conserved candidate virion biogenesis module encoded by IME1s, these findings provide evidence that IME1s produce extracellular virus particles. We hypothesize that that IME1 virions are released without host cell lysis and may have a filamentous morphology. However, because membrane vesicles and other particle types are also abundant in the viral fractions, IME1 morphology will remain unresolved until the genome can be physically linked to a specific particle type. Sequencing of physically separated particle populations, for example following density-gradient fractionation, could establish this link.

**Table 3.** Similarities and differences between IME1 and filamentous bacterial viruses and archaeal budding viruses. *Differences are shown in italic font*.

| **Feature** | **IME1s** | **bacterial inoviruses** | **archaeal budding viruses** |
| --- | --- | --- | --- |
| **pI-like ATPse** | ✓<br>HOG04 | ✓<br>all have pI-like ATPse | ✓<br>Some have pI-like ATPse |
| **Hydrophobic Capsid proteins** | ✓<br>HOG00, HOG01<br>Structural similarity to capsid protein from SSV1 | ✓ | ✓ |
| **Large genome size** | ✓<br>20 - 28kb<br>likely dsDNA | ✗<br>7-8 kb | ✓<br>> 14kb dsDNA |
| **dsDNA genome** | ✓<br>dsDNA | ✗<br>*ssDNA* | ✓<br>dsDNA |
| **Non-lytic** | likely | ✓ | ✓ (varies) |
| **Integrates in host genome** | ✓ | ✓ (varies) | ✓ (varies) |
| **Filamentous morphology** | likely | ✓ | ✓ (varies) |

Standard virus classification tools, including VIPtree, VContact2 (27), and VITAP (28), failed to assign IME1s to a recognized virus group, consistent with their classification as a novel lineage. By contrast, the Inovirus-detection pipeline of Roux *et al*. (8) identified the conserved central region of IME1s as a candidate viral sequence. Broader sampling of Dehalococcoidia and other Chloroflexota will be required to define the distribution and evolutionary history of this virus lineage.

Although no reductive dehalogenase genes were identified in IME1s, several elements encode putative accessory proteins with functions not obviously related to core virus biology, including a possible radical-SAM-domain protein in DmVWBC2 (CDS 22; Table S2.13) and a phosphoadenosine phosphosulfate reductase-like protein in DmVIBARAKI (CDS 28; Table S2.19). These putative accessory genes raise the possibility that IME1s influence host physiology beyond virus production. If IME1s establish chronic, non-lytic infections as proposed, prolonged coexistence of the viral and host genomes could provide opportunities for these elements to influence *Dehalococcoides* physiology and evolution.

## Materials and Methods

### Enrichment Cultures

The KB-1 enrichment culture originated from microcosms established in 1996 using aquifer material from a trichloroethene (TCE)–contaminated site in southern Ontario (29). The parent culture (KB-1/TCE-MeOH) has been maintained for decades with ∼100 mg L⁻¹ TCE as the electron acceptor and methanol (MeOH) as the electron donor (30–32). A vinyl chloride–respiring subculture (KB-1/VC-MeOH) was derived from KB-1/TCE-MeOH and maintained with ∼55 mg L⁻¹ VC and MeOH, as previously described (7). All cultures were maintained anaerobically in batch-fed reactors in a defined pre-reduced anaerobic mineral medium (33).

### Separation of viral fraction and DNA extraction

Viral fractions from KB-1/TCE-MeOH and KB-1/VC-MeOH cultures were collected in 2019 using a protocol adapted from Waller *et al.* (21). Briefly, 500 mL of culture was anaerobically sonicated on ice (40 W, 1 s pulses, 5 min) to disrupt flocs and centrifuged (6000 × g, 30 min, 4 °C). The supernatant was filtered through a 0.45 µm PES membrane (Thermo Scientific™, USA), and the filtrate was amended with 1M NaCl and 10% w/v PEG-8000 (MilliporeSigma, USA) on ice, and incubated overnight at 4 °C. Viral particles were pelleted by centrifugation (5250 × g, 1 h, 4 °C), and resuspended in 1 mL SM buffer (50 mM Tris-HCl pH 7.5, 100 mM NaCl, 10 mM MgSO4, 0.01% gelatin). A second virus-fraction preparation was performed in 2020 without the sonication step, and a third preparation was performed in 2026 using sonication as in the first case. Aliquots from each prep (50 µL) were saved for transmission electron microscopy (TEM; see TEM Methods).

The remaining suspensions of the first and third virus-particle preparations were treated with DNase I (20 U; Thermo Scientific™, USA) for 15 min at room temperature, followed by capsid disruption with Proteinase K (4 µL, 20 mg mL⁻¹; Thermo Scientific™, USA) at 55 °C for 30 min. Viral DNA from 2019 prep was extracted using the Phage DNA Isolation Kit (Norgen Biotek, Canada), eluted in 75 µL Elution Buffer B, and stored at −80 °C. DNA from the 2026 virus fraction was extracted using a phenol-chloroform method (See Supplemental Text 5).

### Metavirome sequencing and assembly

DNA extracted from viral fractions of the KB-1/TCE-MeOH and KB-1/VC-MeOH cultures in 2019 were sequenced at the Genome Québec Innovation Sequencing Centre using Illumina NovaSeq 6000 and PacBio Sequel II platforms. Due to low DNA input, the Illumina libraries were PCR-amplified prior to sequencing. Illumina paired-end sequencing (2 × 150 bp; ∼250 bp insert size) generated approximately 140 million reads per culture. PacBio continuous long-read sequencing yielded ∼3 million reads for KB-1/TCE-MeOH and ∼2 million reads for KB-1/VC-MeOH. Illumina reads were quality assessed using FastQC and trimmed with Trimmomatic (34). Trimmed Illumina reads were assembled using MEGAHIT (35). Hybrid assemblies were subsequently generated with hybridSPAdes in SPAdes v 3.14.1 (36) using MEGAHIT v.1.2.9 contigs as untrusted contigs, followed by scaffolding with SSPACE v2.0 (37). Viral and virus-like sequences were identified using VirSorter2 v.2 (38). Other tools including PhaBOX and VIBRANT for identification and classification were also explored (39, 40).

To improve assemblies of IME1 elements, quality-trimmed Illumina reads were deduplicated using dedup.sh from BBMap (https://sourceforge.net/projects/bbmap/), subsampled to 10% using reformat.sh, and assembled alone or in combination with PacBio reads using metaSPAdes (SPAdes v.3.5.15). PacBio reads were also assembled independently using Flye v2.9.6-b1802. Default settings were used for all software. The viral fraction extracted in 2026 was also sequenced using Illumina as described in Supplemental Text 5 and used to confirm the results from the 2019 viromes.

### Mapping of metavirome reads

Because the 2019 Illumina libraries were PCR-amplified prior to sequencing, quality-trimmed reads were deduplicated using the BBMap dedup.sh script to improve abundance estimates and reduce mapping redundancy. This yielded 11.8 million reads for KB-1/TCE-MeOH and 12.2 million reads for KB-1/VC-MeOH. Deduplicated reads were mapped using BBMap to 1) metavirome contigs, 2) *Dehalococcoides mccartyi* the five closed genomes derived from KB-1 metagenomes (each genome mapped independently), 3) VirSorter-identified viral contigs, and 4) previously described *D. mccartyi* tailed virus sequences using default parameters. Reads were also mapped to all identified IME1 sequences using BBMap with the default parameters except that *ambiguous* was set to *random* (ambiguous mappings were assigned randomly). Genome-level coverage was visualized using the ggcoverage package v1.3.0 in R (41). IME1s and *D. mccartyi* virus sequences were also mapped in Geneious Prime v. 2025 to enable fine-scale parameter adjustment and visual inspection; mappings required ≤5% mismatches per read and inclusion of only properly paired reads. PacBio CCS reads were mapped to genomes using minimap2 in Geneious Prime v. 2026. Deduplicated reads were taxonomically classified using Kaiju v1.9.0 with the NCBI *nr* database, as implemented in the KBase platform (42).

### Bioinformatics analysis of IME1

Initially, twenty-four available genomes from the family *Dehalococcoidia* as of 2019 were screened for the presence of IME1 elements. The IME1 elements were identified based on the following criteria: (i) chromosomal integration downstream of a tRNA or pseudo-tRNA gene; (ii) the first protein-coding sequence encodes a tyrosine recombinase of the XerCD family; (iii) presence of a large coding sequence that may contain a putative metallophosphoesterase domain; (iv) presence of a bifunctional DNA primase–helicase; (v) a terminal coding sequence encoding a Cro/C1-type helix–turn–helix (HTH) repressor; and (vi) an overall element size of ∼23.0 ± 2.0 kb. Subsequently, additional Dehalococcoidia genomes published between 2019 and 2025 were screened for IME1s.

Protein-coding sequences (CDSs) of IME1 elements were predicted using DRAM (43). Additional CDSs were identified by manual curation in Geneious Prime v.2025 following pairwise comparisons of IME1 sequences. The resulting CDSs were clustered into hierarchical orthologous groups (HOGs) using OrthoFinder v3.0.1b1 (44). The HOGs were labelled HOG00 – HOG56; note that this designation is specific for this dataset. Predicted proteins were annotated using BLASTP searches against the NCBI *nr* database (45) and InterProScan, as implemented in Geneious Prime v2025.

Representative proteins from each HOG were further analysed using 1: HHPred (46), 2: PSI-BLAST (47), and/or 3: AlphaFold 3 via the AlphaFold Server (48), and predicted structures were compared using Foldseek (49) against BFVD 2023-2 (50), AlphaFold/Swiss-Prot v6, and PDB100 (20240101).

IME1 nucleotide sequences were aligned using minimap2 v.2.28. The OrthoFinder species tree, based on concatenated shared proteins, was used to order IME1 elements for comparative analyses, which were visualized using gggenomes v1.0.1 in R.

### Phylogenetic analyses

For the HOG04 (Zot-homolog) phylogeny, the structure of the DmVKBm-1 CDS 6 encoded protein was predicted by Alphafold 3. The most similar structures in BFVD 2023-2, MGNIFY_ESM30 and PDB were obtained using FoldSeek. The structures were aligned using FoldMason (51). The alignment was imported to Geneious Prime 2025 and positions with > 50% gaps were removed. Phylogenetic trees were constructed using IQ-TREE 3 with the BLOSUM62+F+I+G4 model and 1000 bootstrap replicates and RAxML v. 8.2.11 in Geneious Prime 2025 with the BLOSUM62 GAMMA model and 100 bootstrap replicates.

### Transmission electron microscopy (TEM) of the viral fractions of KB-1 cultures

Viral suspensions from KB-1 subcultures were applied to Parafilm™, and copper transmission electron microscopy (TEM) grids were floated on droplets for 2 min. Grids were negatively stained with 2% (w/v) uranyl acetate for 2 min, excess stain was removed by blotting with filter paper, and grids were air-dried at room temperature for 10 min. Imaging was performed at the Nanoscale Bioimaging Facility (The Hospital for Sick Children, Toronto, Canada) using a Tecnai T20 transmission electron microscope operated at 200 kV.

### Nucleotide sequence accession numbers

The 2019 TCE and VC metavirome reads and assemblies were deposited to NCBI under the BioSample accession numbers SAMN19349486 and SAMN19349487, respectively, and the BioProject accession: PRJNA732886. The 2026 virome reads were deposited to NCBI under the same BioProject number as BioSample SAMN62356191.

## Acknowledgements

The authors wish to thank Dr. Ali Darbandi at the Nanoscale Bioimaging Facility, The Hospital for Sick Children, Toronto, Canada for assistance with microscopy. The authors also wish to thank Diane Bona and Landon Getz from the Maxwell lab for training and help with viral-fraction isolation. Funding was provided by the Government of Canada through Genome Canada and the Ontario Genomics Institute (no. 2009-OGI-ABC-1405 to E.A.E.), Natural Sciences and Engineering Research Council (NSERC) of Canada (Student Scholarships to O.M. and Discovery grant funding to E.A.E). Support was also provided by the Government of Ontario through the ORF-GL2 program and the Unites States Department of Defense through the strategic Environmental Research and Development Program (SERDP) under the contract (W912HQ-07-C-0036 project ER-1586).

## Conflict of Interest

Authors declare that they have no conflict of interest.

## Notes

### Competing Interest Statement

The authors have declared no competing interest.

### Summary of Updates

Added Supplemental information that was not transferred over from the Journal where we submitted and who have an option to submit to BioRxiv

