## Supplemental for "A novel virus lineage is abundant in metaviromes from *Dehalococcoides*-containing mixed cultures"

Camilla L. Nesbø[1], Nadia Morson[1], Olivia Molenda[1], Line Lomheim[1], Julien Lossouarn[2], Karen L. Maxwell[3], Elizabeth A. Edwards[1]#

[1] Department of Chemical Engineering and Applied Chemistry, University of Toronto, Toronto, ON, Canada
[2] Université Paris-Saclay, INRAE, AgroParisTech, Micalis Institute, Jouy-en-Josas, France.
[3]Department of Biochemistry, University of Toronto, Toronto, ON, Canada

**TABLE OF CONTENTS (19 pages total)**

**Supplemental Tables (All in one Excel file):**

**Table S1:** Summary of annotations of hierarchical orthologous groups (HOGs).
**Table S2.1 to Table S2.21:** Annotation of each IME1 analysed (***Note: 21 Tables***)
**Table S3:** Results from mapping virome reads to IME1 sequences
**Table S4:** Results from mapping virome reads to known *Dehalococcoides* phage sequences
**Table S5A and S5B:** Assembly statistics for the two viromes. The assemblies are hybrid assemblies using both illumina and PacBio data.
**Table S6:** BLAST analysis of KB-1/TCE-MeOH virome contigs with average sequencing depth > 100.
**Table S7:** BLAST analysis of KB-1/VC-MeOH virome contigs with average sequencing depth > 100.
**Table S8:** VirSorter identified phage and phage-like sequences in KB-1/TCE-MeOH metavirome hybrid assembled contigs.
**Table S9:** VirSorter identified virus and virus-like sequences in KB-1/VC-MeOH metavirome hybrid assembled contigs.
**Table S10:** Contigs from the KB-1/TCE-MeOH and KB-1/VC-MeOH metaviromes that contain rdhA genes and the predicted category.
**Table S11.** Genomes and metagenomes with IME1- or Densinoviridae Sf3-like elements
**Table S12:** Additional *Dehalococcoides* genomes with IME1 elements.

**Supplemental Texts: (pages 3-5)**

**Supplemental Text S1:** Naming convention for *Dehalococcoides* cMAGs.
**Supplemental Text S2:** Identification of viral sequences using Virsorter2 and VIBRANT.
**Supplemental Text S3:** Transmission electron micrography (**TEM)**
**Supplemental Text S4:** Mapping metavirome reads
**Supplemental Text S5:** DNA isolation, equencing and assembly of the 2026 TCE-virome.

**Supplemental Figures: (pages 6-19)**

**Figure S1:** Maximum-Likelihood phylogenetic tree constructed HOG04 (the pI-like protein) homologs obtained using FoldSeek and HHpred
**Figure S2:** Maximum-Likelihood phylogenetic tree of HOG02, the tyrosine recombinase.
**Figure S3:** Mapping deduplicated metavirome reads from the TCE and VC cultures (Green and Yellow Y axes) to other *D. mccartyi* genomes, including KBTCE3 genome (panel A), KBTCE1 genome (panel B) and KBVC2 genome (panel C).
**Figure S4:** Mapping metavirome reads generated in 2026 from the KB1-TCE culture (TCE 2026; green) to the *D. mccartyi* KBTCE2 (A) and *D. mccartyi* KBVC1 (B) genomes.
**Figure S5:** tBLASTx comparison of the two IME1 full-length IME1 contigs assembled from metaviromes to the previously identified IME1 called DmVKBVC1 from the *D. mccartyi* VC1 genome.
**Figure S6:** Negatively stained transmission electron microscopy of the KB-1/TCE-MeOH and KB-1/VC-MeOH virus fractions.
**Figure S7:** Negatively stained transmission electron microscopy of the KB-1/TCE-MeOH and KB-1/VC-MeOH phage fractions and whole culture controls.
**Figure S8:** Analysis of a selection of the IME1 sequences generated using VIP-tree.
**Figure S9** Results from VIP-tree (Nishimura *et al.*, 2017) analysis of the IME1 sequences and other contigs classified as Densinoviridae by (Roux *et al.*, 2019) or identified in PSI-BLAST searches using the Zot-like protein (HOG04).

#### SUPPLEMENTAL TEXTS:

##### Supplemental Text S1. Naming convention for *Dehalococcoides* cMAGs.

The naming convention developed for the complete, circular metagenome-assembled genomes (cMAG) identifies culture (KB-1) electron acceptor (in this case vinyl chloride or VC) and relative abundance (number 1 for highest abundance, 2 for next and so on). Thus, the *D. mccartyi* strains from culture KB-1/VC-H2 are named strains KBVC1 and KBVC2. Three *D. mccartyi* genomes containing the vinyl chloride reductase bvcA were closed from the KB-1/1,2-DCA enrichment culture referred to as strains KBDCA1, KBDCA2 and KBDCA3. A *D. mccartyi* complete genome containing a *vcrA* gene was assembled from KB-1/TCE-MeOH culture, named strain KBTCE1. Additionally, two genomes containing trichloroethene reductase *tceA* genes were assembled from the KB-1/TCE-MeOH culture, named strains KBTCE2 and KBTCE3.

See Figure 1 in Molenda et al, 2019.

Molenda O, Tang S, Lomheim L, Gautam VK, Lemak S, Yakunin AF, Maxwell KL, Edwards EA. 2019. Extrachromosomal circular elements targeted by CRISPR-Cas in *Dehalococcoides mccartyi* are linked to mobilization of reductive dehalogenase genes. ISME J 13:24–38.

##### Supplemental Text S2. Identification of viral sequences using Virsorter2 and VIBRANT.

Virome assemblies were analysed using VirSorter2 (Tables S8 and S9) and VIBRANT. In the TCE-metavirome, VirSorter2 identified viruses predicted to infect *Dehalococcoides*, *Geobacter*, and *Acetobacterium*, as well as viruses with unknown hosts, were detected, but all occurred at low abundance (read depth in the range 2-700; Table S8). A greater number of viral contigs were identified in the VC- metavirome, consistent with the larger size of this assembly (Table S9). These included viruses predicted to infect *Dehalococcoides*, *Acetobacterium*, *Methanothrix*, *Methanoregula*, and viruses with unknown hosts; however, all were present at low abundance (read depth in the range 2-700x; Table S9). Very similar results were obtained by VIBRANT. The majority of the viral contigs identified by VIBRANT not identified by Virsorter2 had coverage in the range 3 - 50. Three VC-virome contigs with > 100x coverage were low-quality predictions and had matches in *Dehalococcoides* genomes and were more likely genomic regions. One TCE-virome contig > 100x coverage identified by VIBRANT corresponded to a partial IME1.

##### **Supplemental Text S3.** Transmision electron micrography (TEM)

Transmision electron micrographs (TEMs) from the virus-fraction preparation that did not involve sonication are shown in Figures S3 and S4. As evidence of the effectiveness of the virome preparation, numerous virus-like particles (VLPs) were observed (Figure S3), demonstrating the presence of different viral lineages in the KB-1 microbial community. Consistent with bioinformatic analyses of preparations that included the sonication step, the most abundant structures were vesicles and filamentous structures (Figure S4 J and K). In addition, consistent with previously identified *Dehalococcoides* proviruses (Johnson *et al.*, 2008; Waller *et* *al.*, 2012), siphovirus-like particles were detected in the KB-1 viral fractions (Figure S3A, S3C and 3H). Other tailed viruses might be present as exemplified by putative podovirus-like particles (Figure 3G). Different types of lemon-shaped particles resembling the archaeoviruses of the family *Fuselloviridae* were also observed (Figure 3F and 3I), as well as VLPs with other morphotypes (Figure S3B and S3D).

Johnson DR,Brodie EL, Hubbard AE, Andersen GL, Zinder SH, Alvarez-Cohen L 2008. Temporal Transcriptomic Microarray Analysis of "*Dehalococcoides ethenogenes*" Strain 195 during the Transition into Stationary Phase. Appl Environ Microbiol 74(9): 2864 - 2872.

Waller AS, Hug LA, Mo K, Radford DR, Maxwell KL, Edwards EA. 2012. Transcriptional Analysis of a *Dehalococcoides*-Containing Microbial Consortium Reveals Prophage Activation. Appl Environ Microbiol 78:1178–1186.

#### Supplemental Text S4. Mapping metavirome reads

Mapping the metavirome reads to the *Dehalococcoides* genomes did show above-background abundance of the *VcrA*-IME, though at much lower levels than those observed for IME1s. When mapped to the *D. mccartyi* KBTCE1 and *D. mccartyi* KBVC2 genomes (CP019999, CP019969), the *VcrA*-linked IME region showed ~6–10x higher coverage than background in the TCE-metavirome and ~3–5x higher coverage in the VC-metavirome (Figure S1, Table 2). A similar pattern was observed when viromes were mapped to the *D. mccartyi* KBVC1 genome (CP019968), where the same region showed ~2x higher coverage compared to background levels when mapping TCE-virome reads (CP019968; Figure 1, Table 2). This could indicate that the *VcrA*-IME may also be packaged in virions. However, Molenda et al. (2019) found that this IME occurs as a circular episomal replicon in metagenomes. It is therefore more likely that this DNA is part of the background DNA originating from the *Dehaloccoides* cells that was part of the virus-fraction. We did not observe above-background coverage of other *rdh*-gene regions KB1 *D. mccartyi* genomes, consistent with the low coverage of *rdh*-gene-containing contigs in the assembled metaviromes (Table S10).

Molenda O, Tang S, Lomheim L, Gautam VK, Lemak S, Yakunin AF, Maxwell KL, Edwards EA. 2019. Extrachromosomal circular elements targeted by CRISPR-Cas in *Dehalococcoides mccartyi* are linked to mobilization of reductive dehalogenase genes. ISME J 13:24–38.

#### Supplemental Text S5. DNA isolation, sequencing and assembly of the 2026 TCE-virome.

**Sample preparation:** Viral fraction from of the KB/TCE-MeOH culture was prepared again in 2026 using a similar PEG-precipitation method as in 2019 except that the culture supernatant was filtered through a 0.2 µm filter was used instead of a 0.45 µm filter. Specifically, 1 L of culture was sonicated on ice to disrupt flocs (aiming for 40 W, amplitude 20, tot. energy 21,224 J, 1 s pulses, 5 min) and centrifuged (6000 × g, 30 min, 4 °C). The supernatant was filtered through a 0.2 µm PES membrane (Thermo Scientific™, USA).

**PEG precipitation:** Solid NaCl was added to the filtrate to a final concentration of 1M (salt dissolved, left on ice for 1 hour). Next, 10% w/v PEG-8000 (MilliporeSigma, USA) was added to the filtrate on ice (and allowed to dissolve), and the sample was incubated overnight at 4 °C. Viral particles were pelleted by centrifugation (11000 × g, 1 h, 4 °C), and the pellet was resuspended in 6 mL SM buffer. An aliquot (100 µL) was reserved for microscopy. PEG and cell debris was extracted from the phage suspension by adding an equal volume of chloroform, vortexing for 30s, and the organic and aqueous phases were separated by centrifugation (5000 x g, 15 min, 4°C). This extraction was repeated for the aqueous fraction one more time. The final aqueous sample portion, containing the phage particles, had a volume of 4.5 mL.

**DNA extraction, precipitation and cleanup:** The 4.5 mL phage sample was split into two tubes with 2.25 mL sample each. DNA extraction was done for each tube, using a phenol chloroform method as outlined below. For each tube, 11.25 µl of DNaseI and RNase (10 mg/ml stock) was added and the sample was incubated at 37°C for 1 hour. Next, 5.6 µl of Proteinase K (20 mg/ml stock), 112.5 µl 10% SDS stock (0.5% final concentration), and 90 µl of 0.5 M EDTA pH 8.0 (20 mM final) was added to the tube. The sample was mixed and incubated at 60°C for 1 hour before a 1/10 volume of a 3M sodium acetate solution was added. After allowing the sample to cool to room temperature, an equal volume of cold phenol:chloroform:isoamyl alcohol (30:29:1) was added, and the tube was inverted several times before centrifugation (3000 x g, 5 min, room temp). The supernatant containing the DNA was transferred with a wide-bore pipette tip to a new microfuge tube. An equal volume of phenol:choroform:isoamyl alcohol (30:29:1) was added, the sample vortexed, centrifuged and the supernatant was transferred as before.

The precipitation and cleanup steps described below, were done for each of the two tubes with extracted samples. Isopropanol (0.6 volume) was added before the sample was inverted to mix and centrifuged (max speed, 10 min). The supernatant was carefully removed and the tube with DNA precipitate was filled halfway with 70% ethanol, centrifuged (max speed, 5 min), and as much ethanol as possible was removed without disturbing the pellet. The tube was left open on bench for 15-30 minutes to let ethanol disperse. The final DNA sample was then dissolved in $dH_2O$ (30 µL) and the DNA extract was run on an agarose gel to ensure high MW DNA and pure samples. Finally, the DNA concentration was quantified using Nanodrop and Qubit.

**DNA sequencing and mapping:** The DNA was sequenced at Plasmidsaurus (https://app.plasmidsaurus.com) using a transposase-mediated library preparation chemistry and a 2 x 150 pair-end read configuration, sequenced on an Illumina platform (NovaSeq X Plus) which gave a total of 19,867,550 reads. Mapping of the reads revealed the same pattern as observed in the 2019 virome, with the majority of the reads mapping to IME1 elements (Figure S2). Among megahit assembled contigs, the 12 contigs with > 2000x read depth (2021 – 20,518 x read depth) all correspond to fragments of IME1 elements. Together these fragments cover full IME1 sequences. The fragmentation is likely due to population variation and/or the sonication step used during the virus fraction preparation.

#### Supplemental Figures

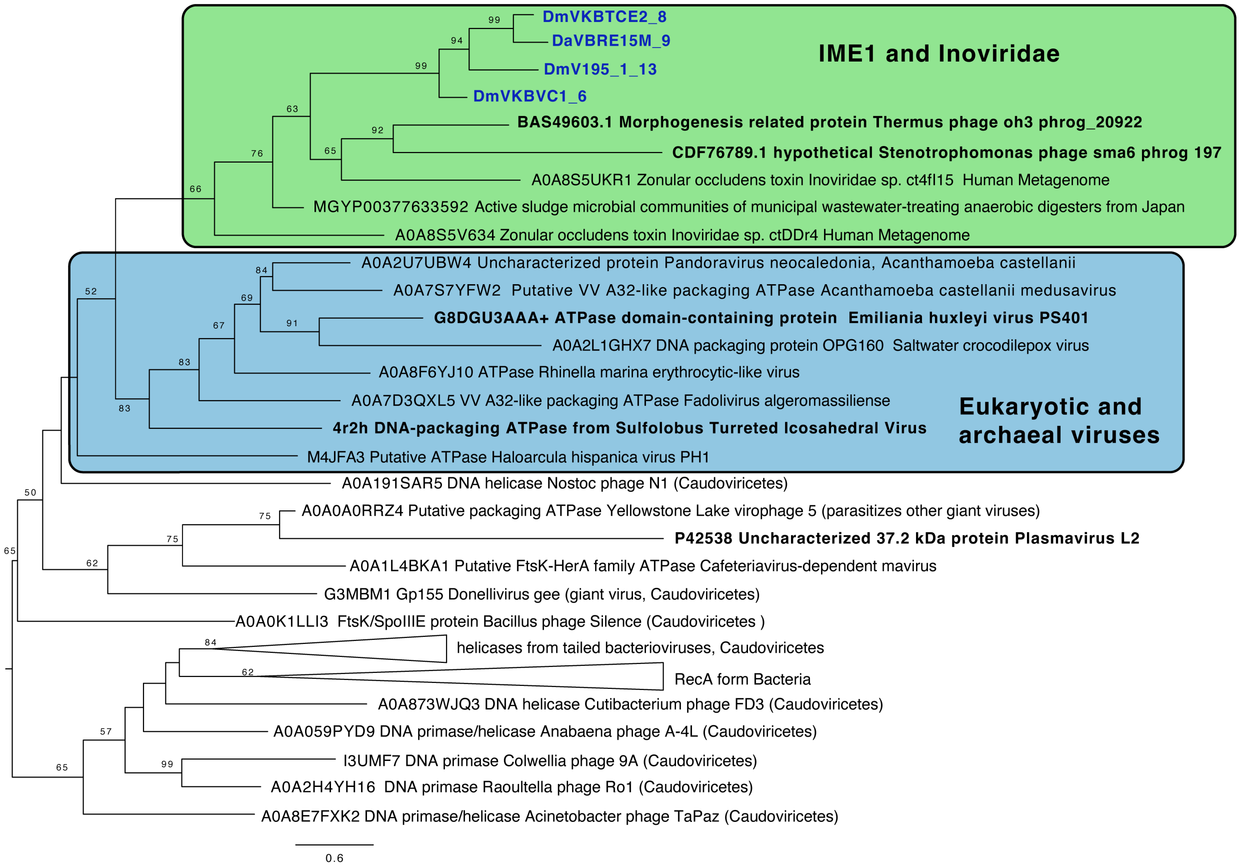


**Figure S1** Maximum-Likelihood phylogenetic tree constructed HOG04 (the pI-like protein) homologs obtained using FoldSeek and HHpred (Van Kempen *et al.*, 2024).
The structure of the DmVKBVC1 CDS 6 encoded protein was predicted by Alphafold 3. A selection of the structures in BFVD 2023-2, focusing on characterized viruses, were obtained using FoldSeek. Structures of representative sequences from phrog 197 and 20922 as well as additional IME1 HOG04 homologs were also predicted in Alphafold 3. The structures were then aligned using FoldMason (Gilchrist et al., 2024), and the alignment was imported to Geneious Prime 2026 and positions with > 50% gaps were removed. The phylogenetic tree was constructed using IQ-TREE 3 with the BLOSUM62+F+I+G4 model and 1000 bootstrap replicates. The topology of a tree estimated by RAxML was the same, however, the support was low for all branches. Sequences in bold represent proteins from characterized viruses known to egress via budding. The green box encloses inoviruses and IME1 while the blue box encloses eukaryotic and archaeal viruses. The taxonomy of proteins from tailed bacterial viruses from the class Caudoviricetes are also indicated.

Van Kempen M, Kim Tumescheit C, Mirdita M, Lee J, Gilchrist CLM, Söding J, Steinegger M. 2024. Fast and accurate protein structure search with Foldseek. Nat Biotechnol 42:243–246.

Gilchrist CLM, Mirdita M, Steinegger M. 2026. Multiple Protein Structure Alignment at Scale with FoldMason. Science 391:485–488

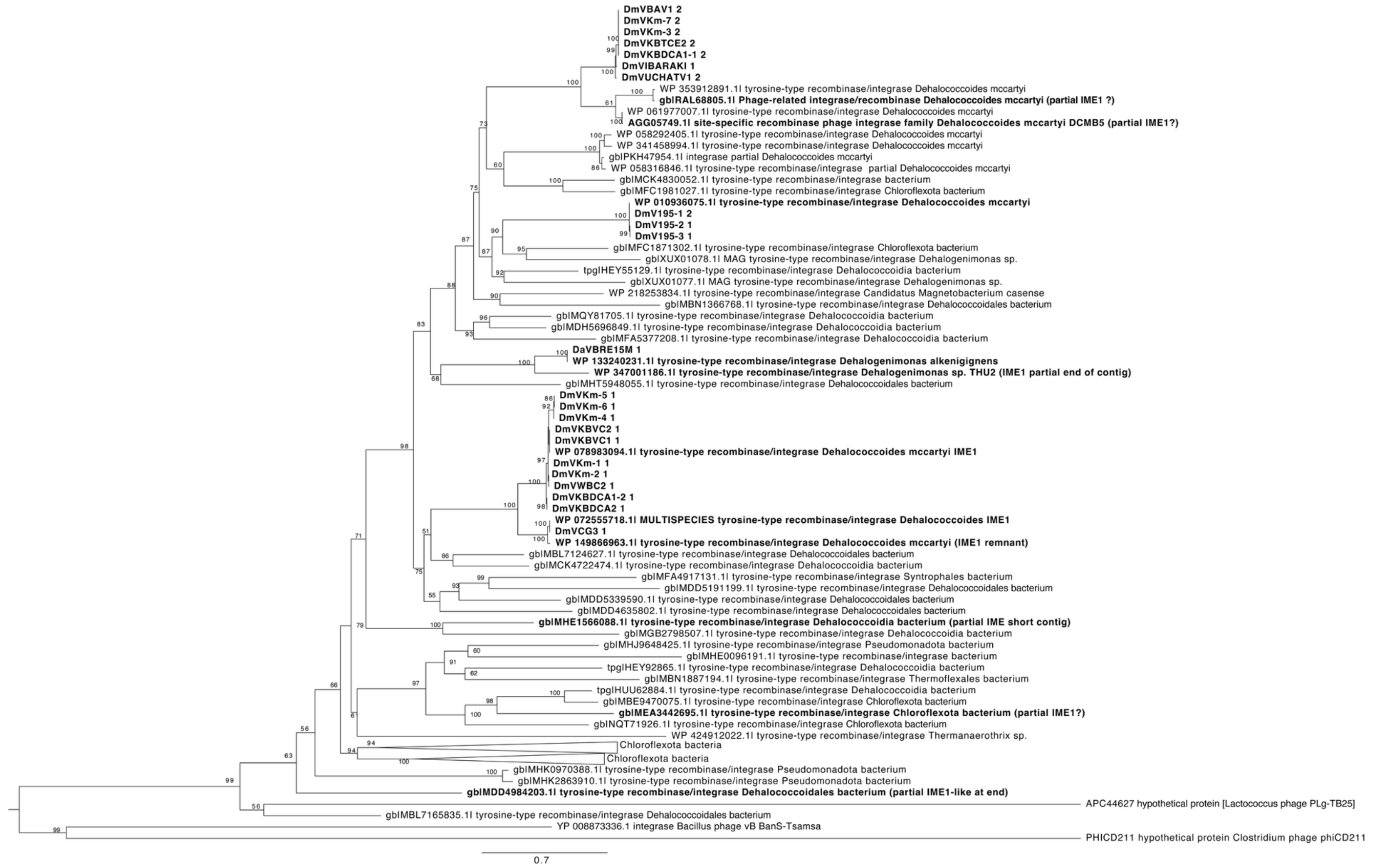


**Figure S2.** Maximum-Likelihood phylogenetic tree of HOG02, the tyrosine recombinase. Homologs of DmVKBVC1 CDS1 were identified using BLASTP against the NCBI ClusteredNR database and the top 100 matches were retrieved and aligned to HOG2 proteins using MAFFT in Geneious Prime v. 2026. The tree was constructed using IQ-TREE 3 with the Q.YEAST+F+R6 model. Numbers on nodes indicate bootstrap support from 1000 replicates. Sequences in bold are from IME1 elements described here or in likely IME1-like elements identified by inspecting the contigs the proteins originated from. Note that the majority of the contigs harbouring the proteins in this analysis were small so IME1-like elements would not be identifiable. Moreover, many of the proteins were encoded by genes at the end of the contig which would also make it hard to associate it with an IME1-like element. APC44627, PHICD211 and YP 008873336.1 were identified as best structural matches in HHpred searches against the Phrog database.

### Figure S3

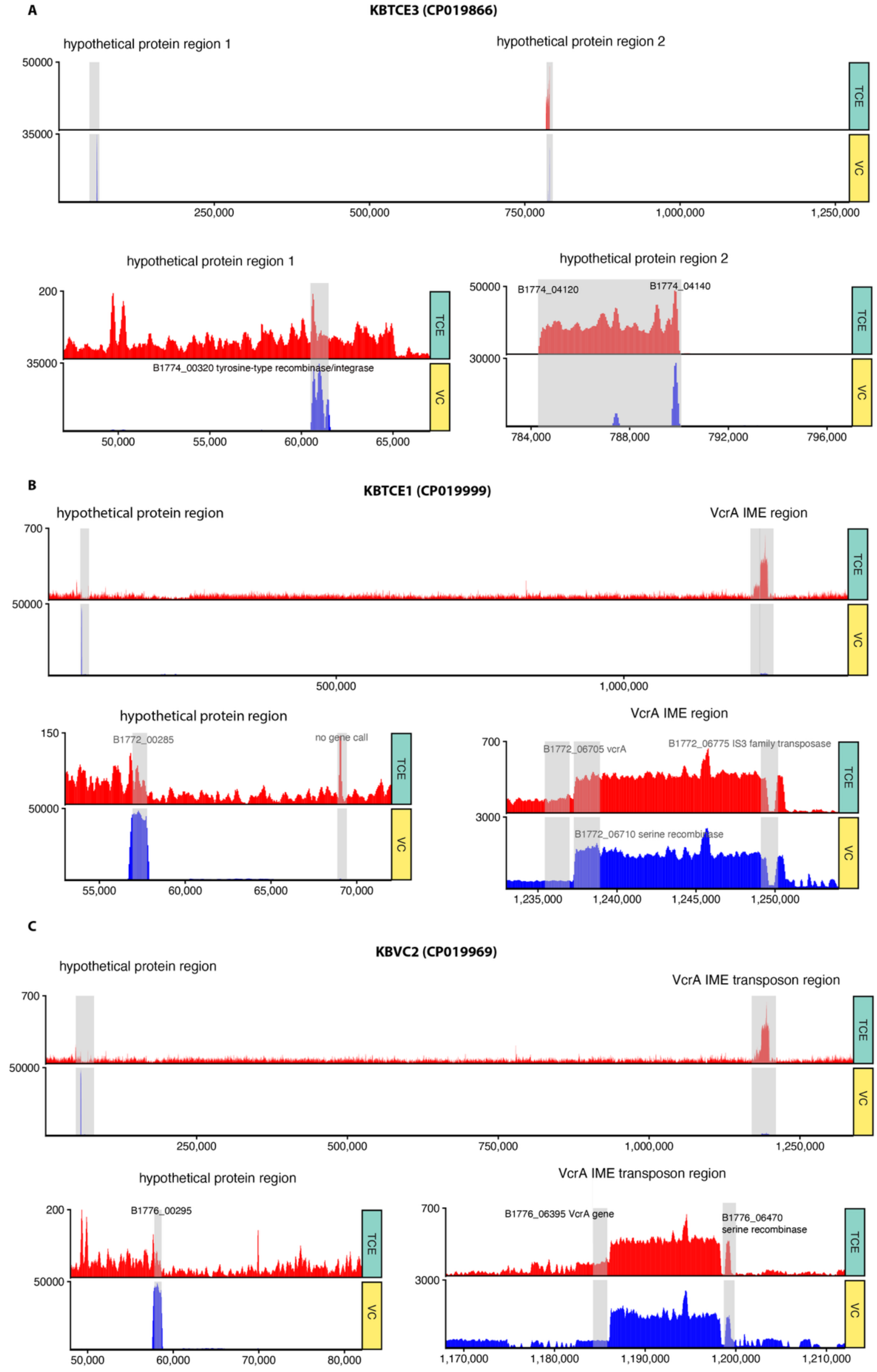
A
KBTCE3 (CP019866)
hypothetical protein region 1
hypothetical protein region 2
50000
35000
TCE
VC
250,000
500,000
750,000
1,000,000
1,250,000
hypothetical protein region 1
200
35000
B1774_00320 tyrosine-type recombinase/integrase
50,000
55,000
60,000
65,000
hypothetical protein region 2
50000
30000
B1774_04120
B1774_04140
784,000
788,000
792,000
796,000
B
KBTCE1 (CP019999)
hypothetical protein region
VcrA IME region
700
50000
500,000
1,000,000
hypothetical protein region
150
50000
B1772_00285
no gene call
55,000
60,000
65,000
70,000
VcrA IME region
700
3000
B1772_06705 vcrA
B1772_06775 IS3 family transposase
B1772_06710 serine recombinase
1,235,000
1,240,000
1,245,000
1,250,000
C
KBVC2 (CP019969)
hypothetical protein region
VcrA IME transposon region
700
50000
250,000
500,000
750,000
1,000,000
1,250,000
hypothetical protein region
200
50000
B1776_00295
50,000
60,000
70,000
80,000
VcrA IME transposon region
700
3000
B1776_06395 VcrA gene
B1776_06470
serine recombinase
1,170,000
1,180,000
1,190,000
1,200,000
1,210,000

**Figure S3** Mapping deduplicated metavirome reads from the TCE and VC cultures (Green and Yellow Y-axis labels) to other *D. mccartyi* genomes, including KBTCE3 genome (panel A), KBTCE1 genome (panel B) and KBVC2 genome (panel C). **Note these genomes do not contain IME1s, unlike those shown in Figure 2**. The upper panel for each genome shows mapping along the full genome sequence, with read depth shown on the Y-axis for the TCE metavirome (TCE - Green) and VC metavirome (VC - Yellow). Regions showing high read recruitment are zoomed in below for each genome. Note that both viromes contain low levels of genomic *D. mccartyi* sequences (Table S5). The average background read-depth for the TCE-virome mapped to the KBTCE3 (CP019866) genome is 103x to the KBTCE1 (CP019999) genome (excluding the VcrA IME region) and is 30x and to the KBVC2 (CP019969) is 30x. The average background read-depth for the VC-virome mapped to the KBTCE3 (CP019866) genome is 188x, to the KBTCE1 (CP019999) genome (excluding the VcrA IME region) is 324x and to the KBVC2 (CP019969) is 321x. **See Table 2 for complete tabulated data.**

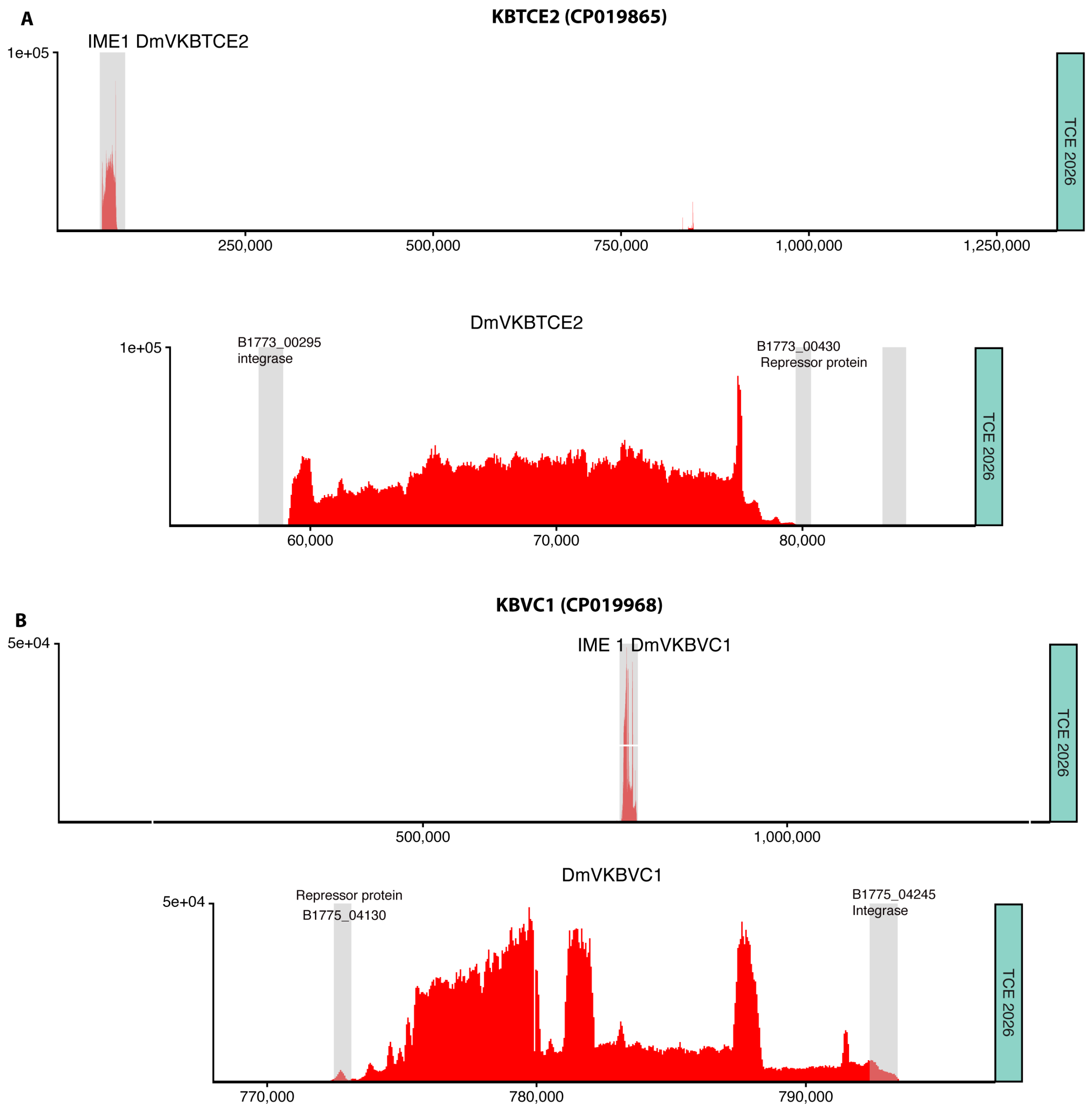


**Figure S4** Mapping metavirome reads generated in 2026 from the KB1-TCE culture (TCE 2026; green) to the *D. mccartyi* KBTCE2 (A) and *D. mccartyi* KBVC1 (B) genomes. The upper panel for each genome shows mapping along the full genome sequence, with read depth shown on the Y-axis. The lower panel shows zoomed in mapping in the IME1 regions. The figure was drawn using ggcoverage v. 1.3.0 in R. The average read-depth for these 2 genomes and other KB-1 genomes without IME1 can be found in Table 2.

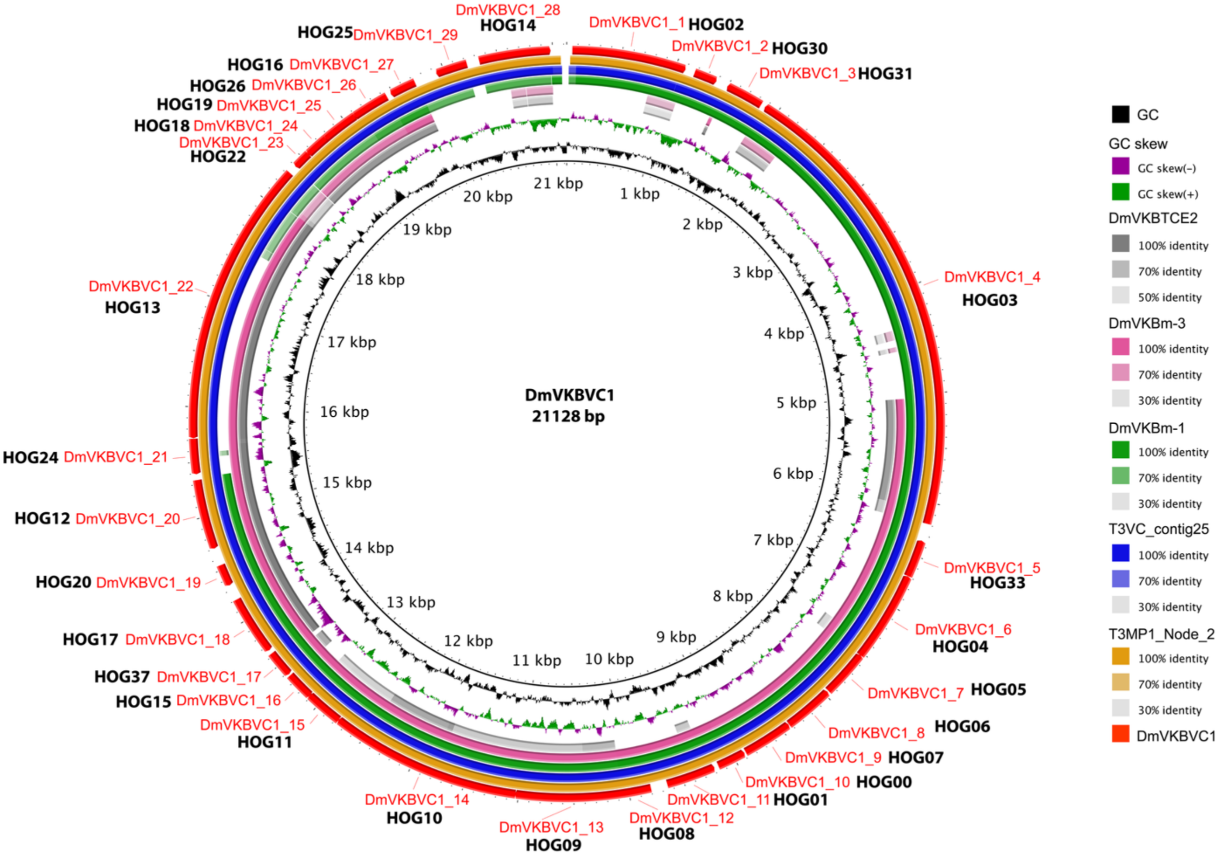


**Figure S5** tBLASTx comparison of the two IME1 full-length IME1 contigs assembled from metaviromes to the previously identified IME1 called DmVKBVC1 from the *D. mccartyi* VC1 genome. Three additional previously identified IME1s from the KB1 cultures are also included. The circles from inner to outer show 1: GC content of DmVKBVC1, 2: GC skew of DmVKBVC1, 3: BLASTN comparison of the DmVKBTCE2 IME1 from the *D. mccartyi* TCE2 genome in grey, 4: BLASTN comparison of the DmVKBm-3 IME1 from the KB1-metagenome in pink, 5: BLASTN comparison of the DmVKBm-1 IME1 from the KB1-metagenome in green, 6: BLASTN comparison of the T3VC_contig25 obtained from the Flye assembly of PacBio reads from the KB-1/VC-MeOH metavirome in blue, 7: BLASTN comparison of the T3MP1 _NODE_2 contig obtained from the hybrid spades assembly of PacBio reads and sub-sampled de-duplicated illumina reads from the KB-1/TCE-MeOH metavirome in orange, 8: CDS in the DmVKBVC1 IME1. The figure was drawn using BRIG v.1.0.0 (Alikhan *et al.*, 2011).

Alikhan, NF., Petty, N.K., Ben Zakour, N.L. *et al.* BLAST Ring Image Generator (BRIG): simple prokaryote genome comparisons. *BMC Genomics* **12**, 402 (2011)

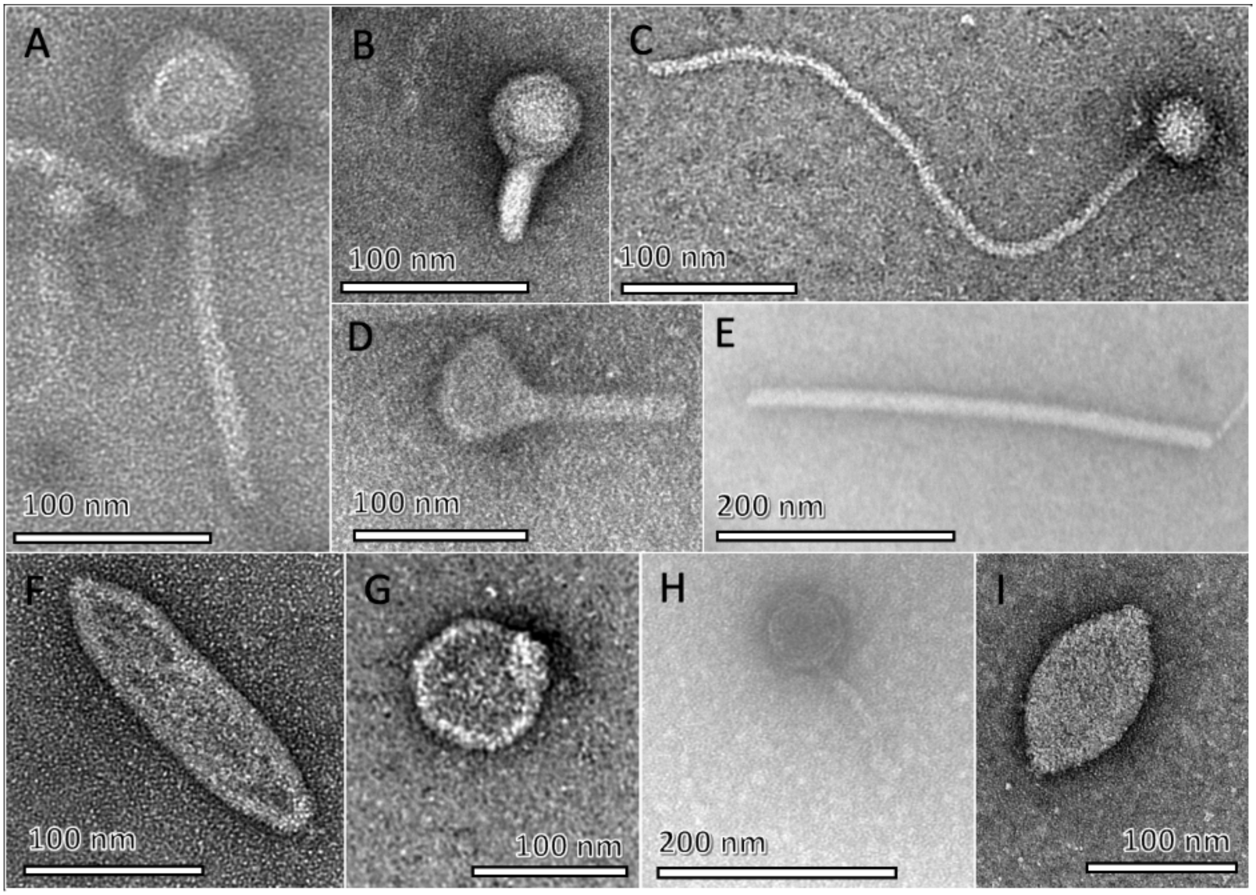


**Figure S6.** Negatively stained transmission electron microscopy of the KB-1/TCE-MeOH viral fraction. All pictures except D were taken from the preparation that did not involve sonication. A variety of virus like particles were observed, including some resembling siphoviruses (A, C and H), podoviruses (G), inoviruses (E), fuselloviruses (F and I), and other possibly archaeal viruses (B, D).

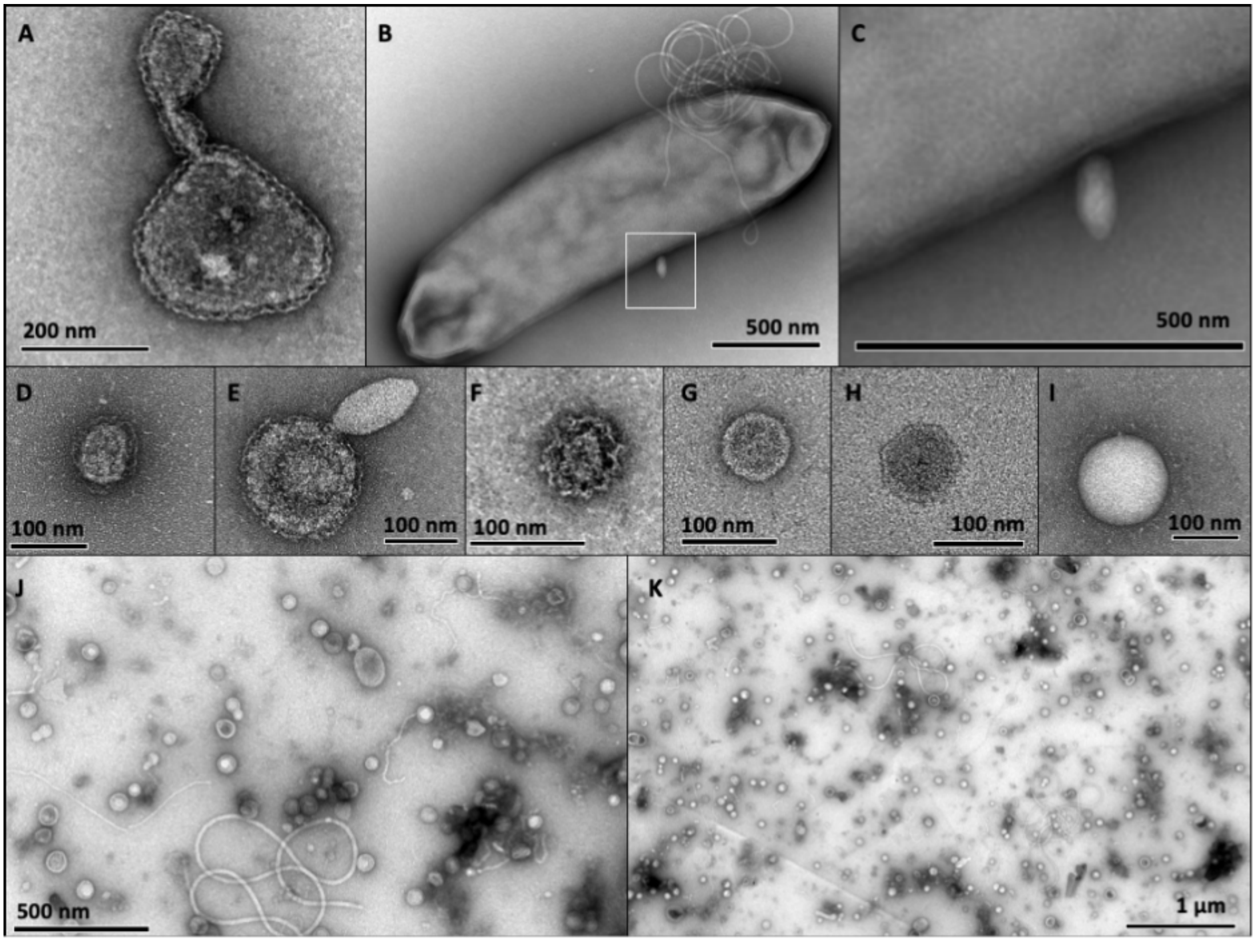


**Figure S7** Negatively stained transmission electron microscopy of the KB-1/TCE-MeOH and KB-1/VC-MeOH phage fractions and whole culture controls. From whole culture controls: (A) a *D. mccartyi* cell undergoing irregular division or blebbing to produce an extracellular structure, and (B) presumably an archaeal cell with a *Fuselloviridae* phage attached (square), and (C) increased magnification for visualization of this phage. From viral fractions: (D), (E) and (F) are S-layered membrane vesicles as visualized by their undulate surface, where (E) may be a *Fuselloviridae* phage attached to a membrane vesicle. (G) and (H) are geometric-shaped particles that may represent encapsulins or tail-less phage-like structures. (I) an unknown spherical structure. (J) and (K) general images from crude virus fractions from KB-1/TCE-MeOH and KB-1/VC-MeOH cultures. Parts of flagella (curly structures) can be seen.

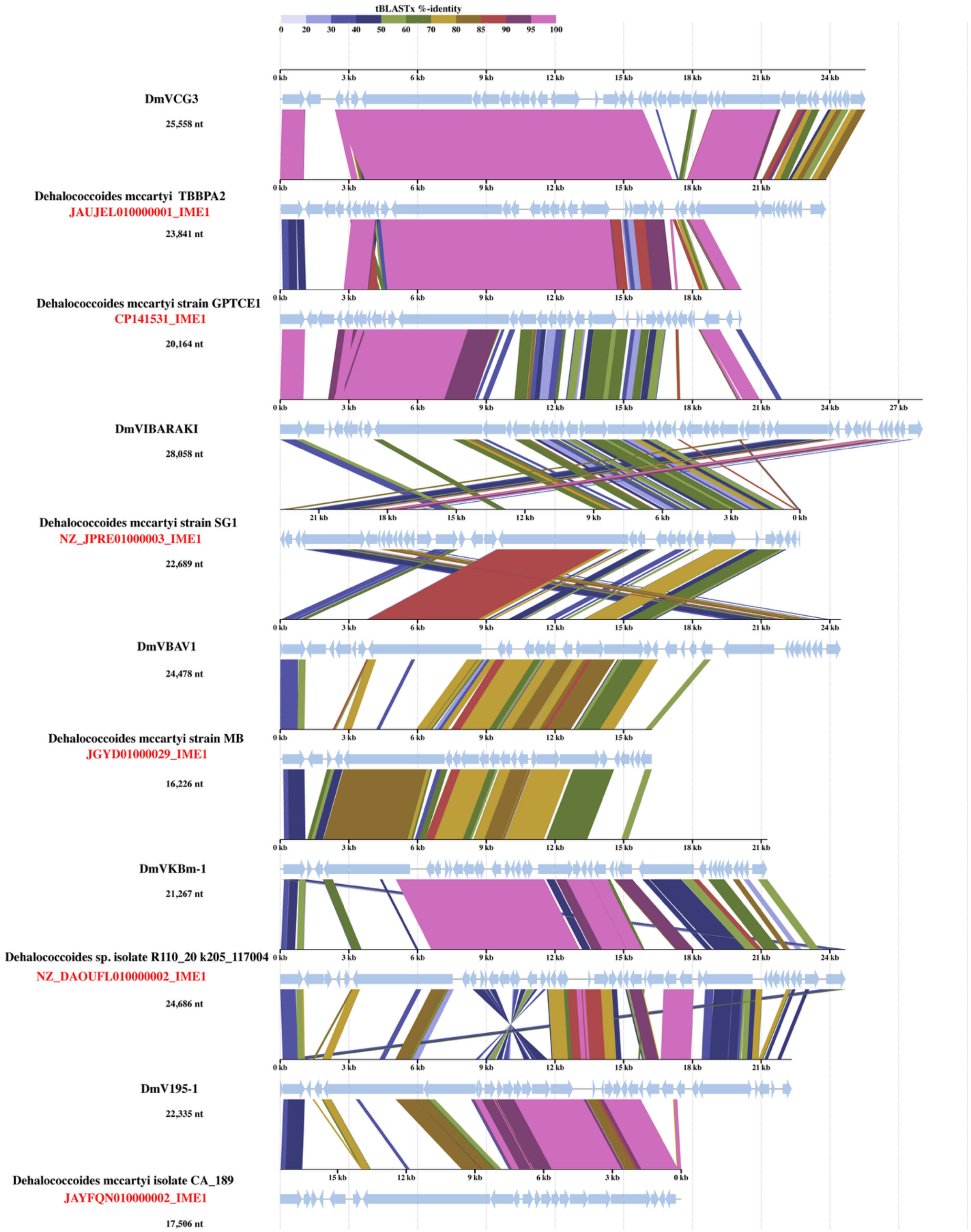
tBLASTx %-identity
0 20 30 40 50 60 70 80 85 90 95 100
DmVCG3
25,558 nt
Dehalococcoides mccartyi TBBPA2
JAUJEL010000001_IME1
23,841 nt
Dehalococcoides mccartyi strain GPTCE1
CP141531_IME1
20,164 nt
DmVIBARAKI
28,058 nt
Dehalococcoides mccartyi strain SG1
NZ_JPRE01000003_IME1
22,689 nt
DmVBAV1
24,478 nt
Dehalococcoides mccartyi strain MB
JGYD01000029_IME1
16,226 nt
DmVKBm-1
21,267 nt
Dehalococcoides sp. isolate R110_20 k205_117004
NZ_DAOUFL010000002_IME1
24,686 nt
DmV195-1
22,335 nt
Dehalococcoides mccartyi isolate CA_189
JAYFQN010000002_IME1
17,506 nt

**Figure S8** Analysis of a selection of the IME1 sequences generated using VIP-tree. The analysis includes a selection of the IME1 sequences analyzed in detail in this study and additional IME1-elements identified in *Dehalococcoides* genomes by PSI-BLAST searches using the pI-like protein (HOG04). The genomes and gene coordinates of these IME1 elements can be found in Table S11. The figure shows the alignment of the sequences generated by VIP-tree (Nishimura *et al.*, 2017). The new IME1 elements are shown in red. The vertical bars indicate similarity detected using tBlastX.

Nishimura Y, Yoshida T, Kuronishi M, Uehara H, Ogata H, Goto S. 2017. ViPTree: the viral proteomic tree server. Bioinformatics 33:2379–2380.

Figure S9

Panel A:

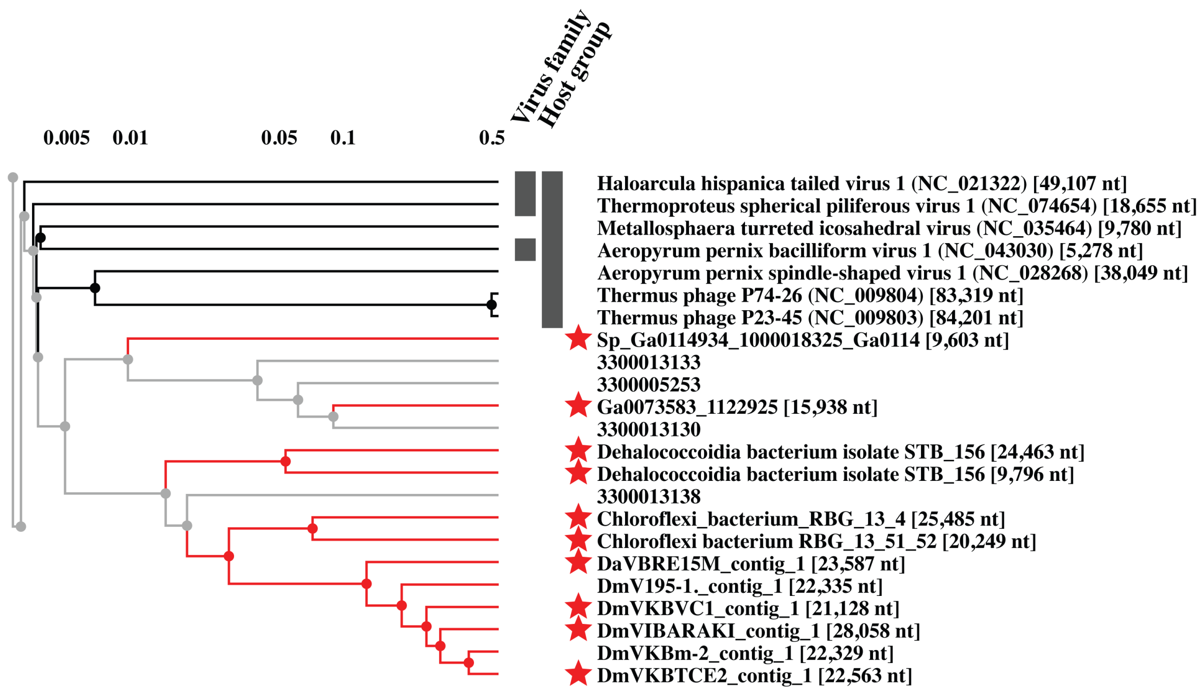

Figure S9

Panel B:

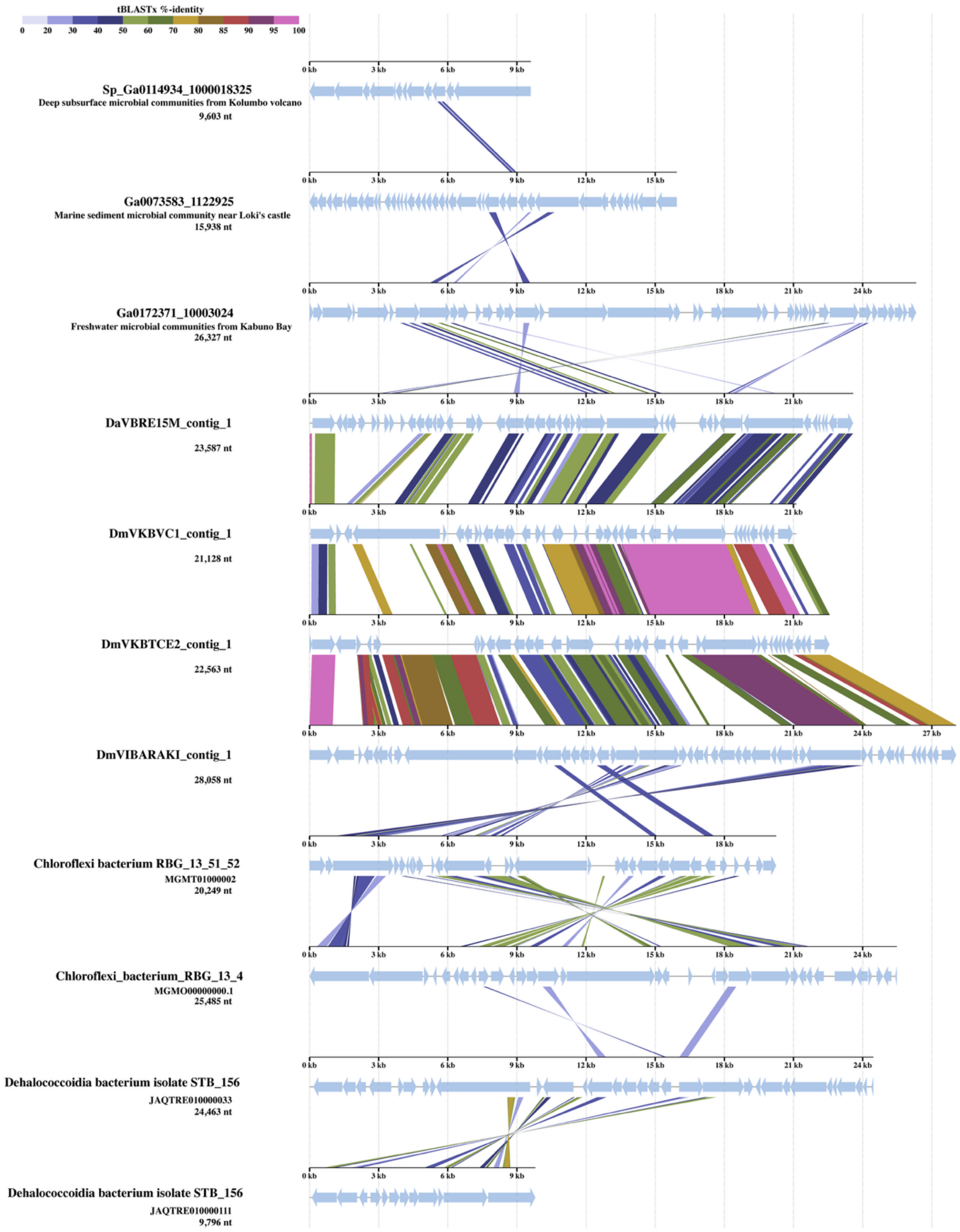

**Figure S9** Results from VIP-tree (Nishimura *et al.*, 2017) analysis of the IME1 sequences and other contigs classified as Densinoviridae by (Roux *et al.*, 2019) or identified in PSI-BLAST searches using the Zot-like protein (HOG04). **A:** The portion of the VIP-tree including the IME1 sequences. **B:** The alignment of the sequences with a red star generated by VIP-tree. The vertical bars indicate similarity detected using tBlastX. Only contigs with alienable regions according to VIP-tree are shown.

Nishimura Y, Yoshida T, Kuronishi M, Uehara H, Ogata H, Goto S. 2017. ViPTree: the viral proteomic tree server. Bioinformatics 33:2379–2380.

Roux S, Krupovic M, Daly RA, Borges AL, Nayfach S, Schulz F, Sharrar A, Matheus Carnevali PB, Cheng J-F, Ivanova NN, Bondy-Denomy J, Wrighton KC, Woyke T, Visel A, Kyrpides NC, Eloe-Fadrosh EA. 2019. Cryptic inoviruses revealed as pervasive in bacteria and archaea across Earth's biomes. Nat Microbiol 4:1895–1906.
